# Enhancing the Identification of NHS Ester-Mediated Lysine-Cysteine Cross-Linking via Reduced Trypsin Digestion Time

**DOI:** 10.64898/2026.09.02.748724

**Authors:** Xin-Feng Yang, Ao-Cheng Hou, Qian Sun, Peng-Zhi Mao, Yu-Jiao Qin, Xiao-Ming Zhang, Guang-Can Shao

## Abstract

Chemical cross-linking mass spectrometry (XL-MS) is a powerful technique for elucidating protein structures and interactions, with NHS ester-based cross-linkers being the most widely used. Traditionally, NHS esters are considered highly specific for primary amines; although their reactivity toward cysteine thiols has been reported, it has not been systematically characterized in XL-MS due to the extreme lability of the resulting thioester bonds. Here, we demonstrate that NHS esters efficiently label cysteine residues and form stable lysine-cysteine (K-C) cross-links—species previously deemed labile in XL-MS. To preserve these thioester-based cross-linked sites, we optimized the standard XL-MS workflow and applied it to four model proteins. K-C cross-links constitute 23% to 59% of total detected cross-links, with Cα-Cα distances predominantly ranging from 12 to 40 Å, and 60% satisfying theoretical linker constraints—confirming that K-C cross-links are as reliable as canonical lysine-lysine (K-K) cross-link sites. Cross-software validation and benchmarking against the heterobifunctional K-C cross-linker GMBS further demonstrated that our optimized workflow captures ∼ 40% of GMBS-identified K-C cross-linked sites. Collectively, these findings expand the practical utility of NHS esters and enhance protein structural characterization by enabling simultaneous acquisition of K-K and K-C distance constraints.

**For Table of Contents Only:** 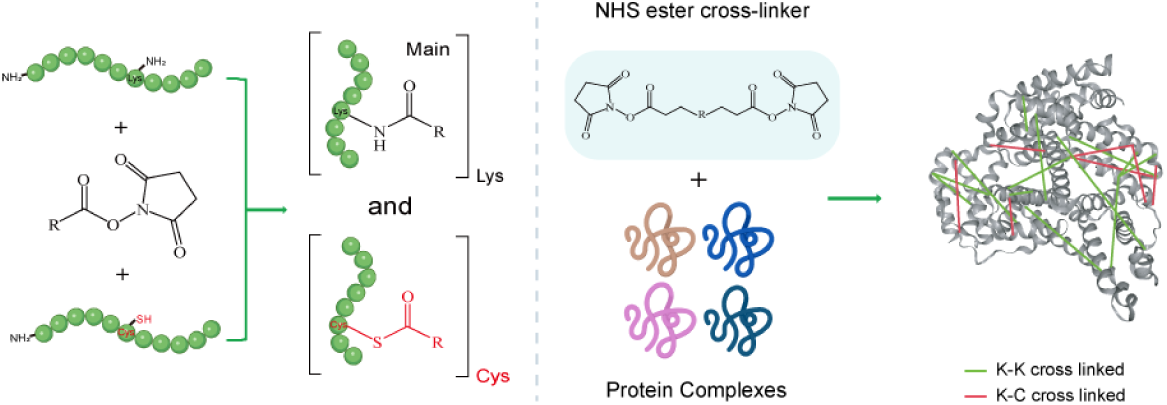

## Introduction

Chemical cross-linking mass spectrometry (XL-MS) has emerged as a robust platform for elucidating protein architectures, characterizing conformational dynamics, and mapping protein-protein interaction networks^1–13^. By covalently tethering proximal amino acid residues, XL-MS generates distance constraints that define upper limits for inter-residue separations, thereby facilitating the structural characterization of complex topologies^14^. The versatility of XL-MS has extended beyond static structural modeling to include in vivo architectural mapping^15–17^, the study of intrinsically disordered regions^18–20^, and the profiling of membrane protein interfaces and cell-surface networks^21, 22^. Furthermore, the method is increasingly employed to probe allosteric transitions and structural rearrangements induced by ligand binding^23, 24^.

Among the diverse chemistries available, N-hydroxysuccinimide (NHS) ester-based cross-linkers are the most prevalent due to their efficient reactivity under mild, physiological conditions^25–27^. Homobifunctional NHS reagents are widely utilized and can be categorized based on their spacer arms: chemically cleavable (e.g., DSP), mass spectrometry (MS)-stable (e.g., DSS, BS3, BS2G), MS-cleavable (e.g., DSSO^28^, DSBU^29^), and enrichable linkers that incorporate phosphate, biotin group or dimethylpiperidinyl group (e.g., PhoX^30^, Leiker^31^, DSBSO^32^, DPST^33^). Additionally, heterobifunctional cross-linkers often pair an NHS ester with a different reactive moiety to expand the range of targetable amino acids, examples include GMBS^34^ (lysine-to-cysteine) and photo-activatable SDA^35, 36^ (lysine-to-any residue). Collectively, these reagents demonstrate excellent performance in protein structure analysis, interface mapping of complexes, and profiling protein-protein interaction networks.

Although NHS esters are widely used because of their high selectivity toward primary amines, a comprehensive understanding of their overall reaction chemistry remains important for expanding their applications in XL-MS and chemical biology^37^. It is well established that NHS esters maintain robust selectivity for primary amines under typical chemoproteomic conditions, particularly in complex proteomes^38, 39^. Reactions of NHS esters with thiols and hydroxyl groups (serine, threonine, and tyrosine) have also long been recognized^40, 41^. However, the structural utility of cysteine reactivity in XL-MS has not been systematically explored.

In this study, we demonstrate that NHS ester reagents can successfully label cysteine residues in synthetic peptides, even in the spatial proximity of lysine. Furthermore, we show that homobifunctional NHS cross-linkers can form lysine-cysteine (K-C) cross-links. These cross-links are conventionally considered labile because the resulting thioester bonds possess high free energy of hydrolysis and pronounced susceptibility to nucleophilic attack^42, 43^. By systematically refining the sample preparation workflow—specifically by utilizing TCEP for reduction, maintaining a near-neutral pH, and minimizing enzymatic digestion time—we significantly improved the recovery and stability of these K-C cross-linked products. The application of this optimized workflow to four model proteins revealed that K-C cross-linked sites constitute 23% to 59% of the total detected cross-links. Validation through structural mapping and searching against entrapment database and entrapment link sites confirmed that these K-C cross-links are as reliable as canonical K-K cross-links. Finally, a comparative analysis indicated that the K-C cross-links generated by DSS captured approximately 40% of the K-C cross-linked sites identified by the classic heterobifunctional K-C cross-linker, GMBS. Together, these findings demonstrate that homobifunctional NHS ester cross-linkers can be repurposed to reliably yield both K-K and K-C distance constraints, thereby substantially enhancing the depth and resolution of protein structural analysis.

## Materials and Methods

### Chemicals and Reagents

Synthetic peptides were provided by Anhui Guoping Pharmaceutical Company. Labeling reagents and the cross-linkers DSSO, DSBU, and GMBS, were purchased from Biodee Pharmaceutical. The cross-linker DSS was purchased from Thermo Fisher Scientific. The following reagents were utilized in the preliminary sample preparation and enzymatic hydrolysis: tris(2-carboxyethyl) phosphine (TCEP), dithiothreitol (DTT), iodoacetamide (IAA), and urea, all obtained from MedChemExpress; trifluoroacetic acid (TFA), formic acid (FA), acetonitrile (ACN), and tris buffer, all sourced from Thermo Fisher Scientific.

### Peptide Labeling and Deprotection

Synthetic peptides and labeling reagents, both pre-solubilized in DMF, were reacted at a 1:1 molar ratio. The reaction was performed in either DMF (supplemented with 2% TEA) or 50 mM HEPES buffer (pH 7.5) at room temperature with constant agitation (800 rpm, 30min). Specifically, cysteine- and lysine-containing peptides, (Ac)MVLCESFHWKVIR and (Ac)DTCIGYEKLAQR, were labeled with NHS-propionyl reagent (*N*-(Propionyloxy) succinimide) at a 1:2 molar ratio in 50 mM HEPES (pH 7.5). For peptides containing N-terminal Boc protection, deprotection was performed following labeling using a cleavage reagent (95:5 TFA: H₂O, v/v) for 20min at room temperature with agitation. The peptide was precipitated twice with pre-chilled diethyl ether and collected by centrifugation (12,000 rpm, 10 min), and finally evaporated to dryness before being re-solubilized in 0.1% (v/v) FA.

### Cross-linking-mediated Assembly of K-C Peptide Pairs

Cross-linking between cysteine- and lysine-containing peptide pairs was conducted in HEPES buffer (50mM pH 7.5). The MS-cleavable cross-linker DSSO and the non-cleavable cross-linker DSS were employed for five distinct peptide pairs, maintaining a 1:1:2 molar ratio (lysine-peptide : cysteine-peptide : cross-linker). Reactions proceeded for 30 min at room temperature under 800 rpm agitation. Following N-terminal Boc deprotection (95% TFA), the samples were dried and reconstituted in 0.1% (v/v) FA for LC-MS/MS analysis.

### Cross-Linking of four Model Proteins

BSA, Cytochrome P450, Lactoferrin, and Lysozyme (30 μg) were individually cross-linked with 1 mM DSS, DSSO, or DSBU in 100 mM HEPES (pH 7.5) for 30 min (1000 rpm). Following quenching (20 mM ammonium bicarbonate, 10 min) and acetone precipitation (-20 °C, ≥2 h), protein pellets were collected by centrifugation, dried, and reconstituted in 8 M urea. Samples were then reduced (5 mM TCEP, 20 min), alkylated (10 mM IAA, 10 min, dark), and diluted to 2 M urea for tryptic digestion (1:20 w/w, 37 °C, 4 h). Digestion was stopped with 1% (v/v) FA. Finally, peptides were SPE-purified, vacuum-dried, and re-solubilized in 0.1% (v/v) FA.

### LC-MS/MS Data Acquisition

LC-MS/MS analysis was conducted on a Vanquish Neo UHPLC system coupled to an Orbitrap Eclipse Tribrid mass spectrometer (Thermo Fisher Scientific). Peptides were loaded onto a 22 cm × 100 μm C18 column, which was packed with 1.9 μm, 120 Å particles (Dr. Maisch GmbH), and were separated at a flow rate of 500 nL min^−1^. The mobile phases A and B consisted of 0.1% FA in water and 0.1% FA in 80% ACN, respectively. The chromatographic gradient was as follows: 0.1 min at 4% B; a linear increase from 4% to 35% B over 53.9 min; and 35% to 99% B over 0.1 min, followed by a re-equilibration period.

MS1 scans were acquired in the Orbitrap (resolution 120,000; *m/z* 375–1500). Precursors with charge states 3+ to 6+ and intensities exceeding 4.0e4 were prioritized for HCD fragmentation (NCE 30%) using an isolation window of 1.6 *m/z*. MS2 spectra were recorded at 15,000 resolution with an AGC target of 7.5e4 and a maximum injection time of 120 ms. Dynamic exclusion was set to 30 s with a 10 ppm tolerance.

### Software and Search Parameters

pLink 3.0.17 and xiSearch 1.8.11 (coupled with xiFDR 2.3.10) were used to analyze the data obtained from cross-linked samples. Cross-linker site settings: pLink, α site: K and n-terminal; β site: K, C and n-terminal. xiSearch, LINKEDAMINOACIDS: K(0), C(0), S(0.2), T(0.2), Y(0.2), nterm(0).

The search parameters for the identification of cross-linked peptide pairs were set as follows: the precursor mass tolerance, 10 ppm; the fragment ion mass tolerance, 20 ppm; variable modification, Carbamidomethyl[C] and Oxidation[M]; peptide length minimum, 6-60 amino acids per chain; peptide mass, 600-6000 Da per chain; enzyme, trypsin; and three missed cleavage sites were allowed. Identifications from both platforms were filtered using a 5% peptide-pair-level FDR, with the “boost” option enabled for xiSearch analysis.

## Results

### Evaluation of the reactivity of NHS Ester Toward Cysteine residue in Synthetic Peptides

Initial observations of mass spectra corresponding to cysteine residue modified by an NHS ester cross-linker (Figure S1A-B) prompted a systematic investigation into the reactivity between NHS esters and the cysteine thiol group. To initiate this investigation, we conducted labeling experiments using synthetic peptides. To isolate the intrinsic reactivity of the cysteine thiol, we employed peptides with N-terminally acetylated (Ac-) or Boc-protected amines to prevent modification of the α-amino group (Figure 1A). These peptides were subsequently reacted with three NHS esters (*N*-(Propionyloxy) succinimide, Biotin 3-sulfo-N-hydroxysuccinimide ester sodium salt, and 2,5-Dioxopyrrolidin-1-yl octanoate) under both organic (DMF with 2% TEA) and physiologically relevant aqueous (50 mM HEPES, pH 7.5) conditions. Following the reaction, the Boc-protecting group was removed with trifluoroacetic acid (TFA) to generate peptides mimicking tryptic digestion products.

**Figure 1.**
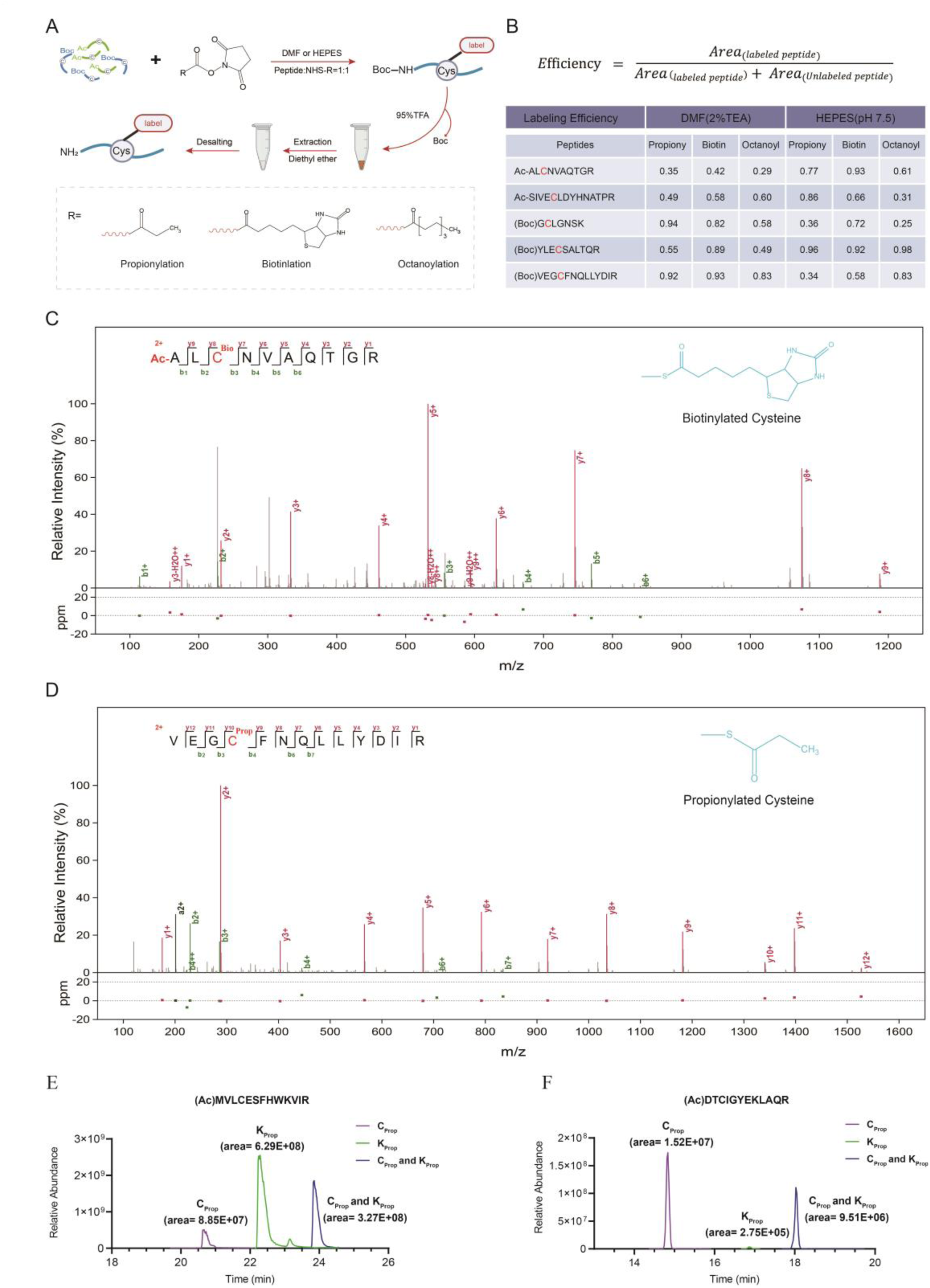
Characterization of NHS ester reactivity toward cysteine thiol groups. (A) Reaction schemes of N-terminally protected, cysteine-containing peptides with three NHS-ester reagents. Acetylated (Ac, green) and Boc-protected (blue) peptides were utilized as substrates. Chemical structures of the varying functional groups (R) are shown below. (B) Labeling efficiency of synthetic peptides under organic (DMF with 2% TEA) and aqueous (50 mM HEPES, pH 7.5) conditions. Efficiency was calculated as the ratio of the labeled peptide peak area to the total peptide peak area (formula provided). Free cysteine residues are highlighted in red. Note: Disulfide-linked peptide peaks were excluded from calculations because their formation consumes free cysteine thiols, directly competing with the labeling reaction. (C-D) Representative MS/MS spectra of biotinylated cysteine peptide (C) and propionylated cysteine peptide (D). (E-F) Competitive propionylation between lysine and cysteine residues. Extracted ion chromatograms (XICs) for (E) (Ac)MVLCESFHWKVIR and (F) (Ac)DTCIGYEKLAQR following reaction with NHS-propionyl. Peak area integration displays the chromatogram of mono-labeled (Cys, purple; Lys, green) and doubly labeled (Lys + Cys, blue) species.

Labeling efficiency was quantified by calculating the ratio of the integrated chromatographic peak area of the labeled peptide to the sum of the peak areas of both unlabeled and labeled species (Figure 1B). The results demonstrated that NHS esters consistently reacted with the cysteine residue across all five synthetic peptides, although the efficiency varied depending on the peptide sequence and buffer system. The median labeling efficiency across 30 independent reactions was 65.8%, with eight reactions exceeding 85% and the highest reaching 98%. High-resolution mass spectra of the biotinylated, propionylated, and octanoylated products provided unambiguous evidence of site-specific cysteine modification (Figure 1C-D, Figure S2A-C). These data confirmed that the cysteine thiol is a viable nucleophilic target for NHS reagents.

Given that the ε-amino group of lysine is the canonical target of NHS esters, we next investigated whether cysteine could be modified in the presence of a proximal lysine residue. We synthesized two peptides containing both cysteine and lysine, as well as other residues with various potentially reactive side chains (Ser, Thr, Tyr, Asp, Glu, His, and Arg). The peptides were then labeled using a sub-stoichiometric amount of NHS-propionyl reagent (reagent-to-peptide ratio of 1:2) to assess competitive reactivity. For the peptide Ac-MVCLESFHWKVIR, the Lys-propionylated species (Kprop) exhibited the highest abundance (peak area= 6.29×10^8^); however, a substantial portion of the detected signal involved cysteine modification (Cprop, 8.85×10^7^; Cprop+Kprop, 3.27×10^8^), indicating that nearly half of the modified peptides were derivatized at the thiol group (Figure 1E). Remarkably, for the peptide Ac-DTCIGYEKLAQR, the cysteine thiol exhibited dominant reactivity. The Cys-propionylated (Cprop, peak area= 1.52×10^7^) and dual-modified (9.51×10^6^) species collectively accounted for 99% of the total peak area, while lysine mono-modification was negligible (2.75×10^5^) (Figure 1F). These competitive labeling experiments definitively establish the cysteine residue as a highly competent target for NHS reagents, with reactivity that can rival or even exceed that of lysine depending on the local sequence context.

### Formation and Identification of Lysine-Cysteine Cross-Links with Homobifunctional NHS Esters

Based on our preliminary findings, we hypothesized that standard homobifunctional NHS ester cross-linkers, such as DSS and DSSO, could generate stable lysine-cysteine (K-C) cross-links.

To test this hypothesis under controlled conditions, we first performed cross-linking experiments using five synthetic peptide pairs. N-terminally Boc-protected peptides, one containing a lysine and the other a cysteine, were reacted with either DSS or DSSO (2:1 cross-linker:peptide ratio). Following the reaction, the Boc groups were removed with TFA to generate peptides mimicking tryptic products for MS analysis (Figure 2A). This experimental design was expected to produce K-K, K-C, and C-C cross-linked species. Quantitative analysis based on chromatographic peak areas revealed that both DSS and DSSO effectively facilitated the formation of K-C cross-links, with signal intensities ranging from 1e6 to 1e9. Although the canonical K-K cross-links were the predominant species, the K-C signal was consistently robust. In contrast, the signal intensities for half C-C cross-linked peptides was below the 1e6 threshold, with the exception of one peptide pair (TFEESFQKALR-GCLGNSK) that yielded high-intensity C-C products, likely due to favorable sequence context or ionization efficiency (Figure 2B). The K-C products were rigorously validated by high-resolution MS/MS analysis, which provided site-specific evidence through extensive b- and y-ion series and showed precursor ion isotopic envelopes in excellent agreement with theoretical distributions (Figure 2C-D, Figure S3A-C).

**Figure 2.**
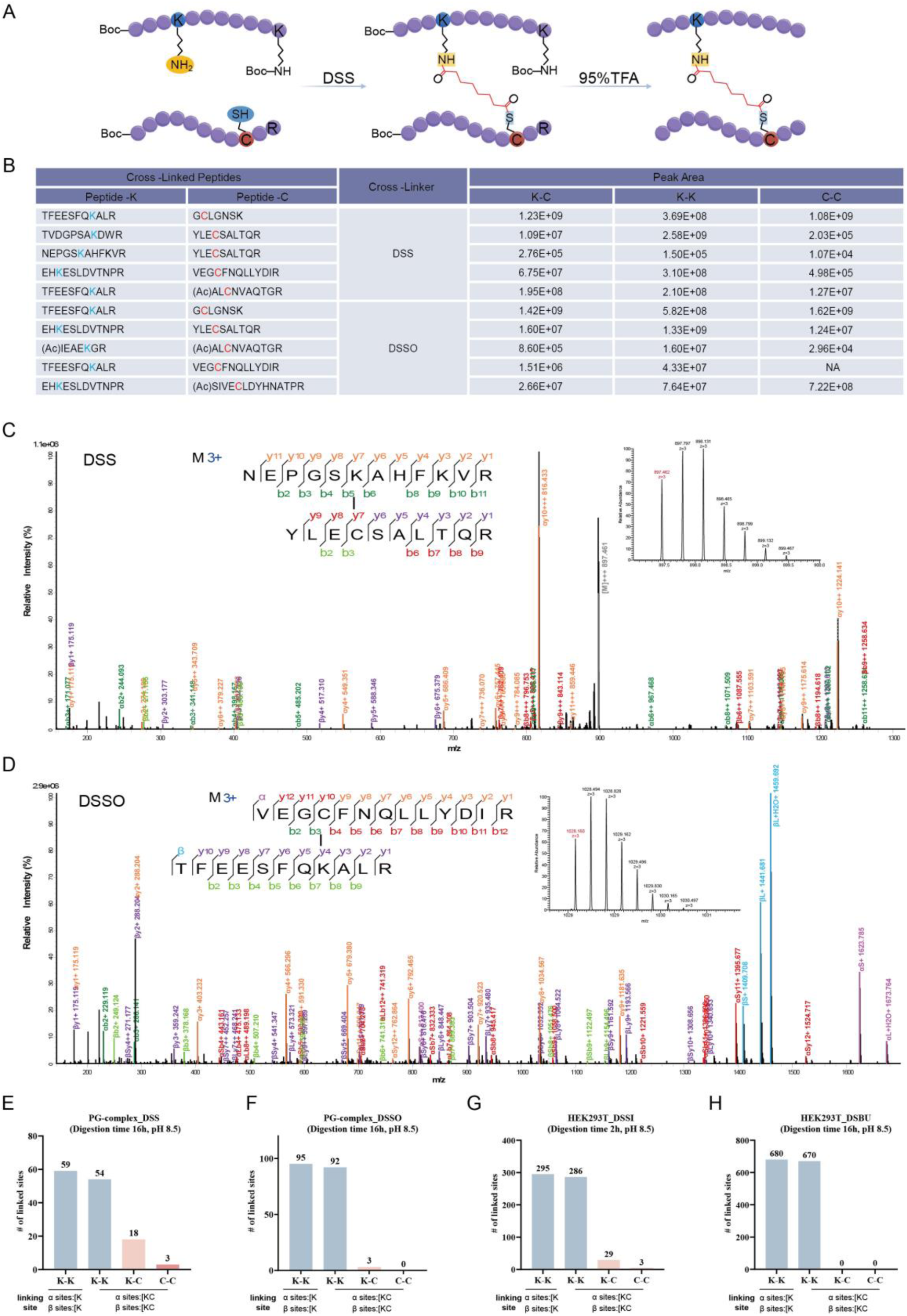
Identification of lysine-cysteine (K-C) cross-links in synthetic peptides and proteomic datasets. (A) Reaction scheme illustrating the formation of cross-links between lysine- and cysteine-containing peptides. (B) Relative proportions of cross-linkage types (K-C, K-K, and C-C) generated using DSS and DSSO for five distinct peptide combinations. Proportions were determined by peak area integration. Reactive Lys (blue) and Cys (red) residues are highlighted within the primary sequences. (C-D) Representative MS/MS spectra of identified K-C cross-linked peptides: (C) NEPGSKAHFKVR-YLECSALTQR (*m/z*=897.462, *z*=3+) and (D) VEGCFNQLLYDIR-TFEESFQKALR (*m/z*=1028.159, *z*=3+). Precursor isotopic clusters are shown in the right insets. (E-H) Re-analysis of public proteomic datasets using carbamidomethylation as a variable modification to identify cysteine-containing cross-links. Results are shown for the PG-complex (PXD075905: E, DSS; F, DSSO) and HEK293T cells (PXD050960: G, DSSI; H, DSBU). Cross-linkage types are color-coded: K-K (blue) and cysteine-containing K-C/C-C (pink/red). Digestion conditions (incubation time and pH) for each dataset are indicated above the respective panels.

In standard XL-MS workflows, cysteine residues are typically reduced and irreversibly alkylated (e.g., with iodoacetamide), and the resulting carbamidomethylation is treated as a fixed modification during database searching. This practice effectively excludes cysteine as a potential cross-linking site. To determine if K-C cross-links exist in complex biological samples, we re-analyzed four public XL-MS datasets by re-defining carbamidomethylation as a variable modification and including cysteine as a reactive residue in our search parameters. K-C cross-links were consistently identified across multiple reagents and digestion protocols (Figure 2E-G). For a hetero-dimeric protein complex (PG-complex), we identified 18 (24%) and 3 (3%) K-C cross-links in DSS and DSSO datasets, respectively. In a DSSI cross-linked HEK293T cell lysate digested for 2 hours with trypsin, we identified 29 (9%) K-C cross-links^44^. Notably, no reliable K-C cross-links were found in a DSBU-linked lysate subjected to a prolonged 16-hour digestion, as compared to the 2-hour digestion control, suggesting potential lability of the thioester bond over extended incubation times (Figure 2H). The number of C-C cross-links across all four datasets was fewer than three, indicating this species is negligible in situ. All identified K-C or C-C cross-links were manually validated, and the representative high-quality spectra are presented (Figure S4A-C). These results demonstrate that NHS ester-mediated K-C cross-links are formed not only on synthetic peptides but also within protein complexes and proteome-wide samples.

### Optimizing the XL-MS Workflow to Preserve Thioester-Based K-C Cross-Links

Given the inherent lability of the thioester bond formed between an NHS ester and a cysteine thiol, we systematically evaluated the standard XL-MS workflow (Figure 3A) to identify and mitigate steps contributing to K-C cross-link loss. We first monitored the stability of DSS-linked products from a synthetic peptide pair (TVDGPSAKDWR and GCLGNSK) throughout a typical urea-based sample preparation protocol. Aliquots were taken after reduction, alkylation, and LC-MS analysis after tryptic digestion for 2, 4, and 16 hours. The results revealed that the use of DTT as a reducing agent resulted in a significant loss of K-C cross-links compared to TCEP. Furthermore, K-C cross-links exhibited time-dependent loss throughout the workflow, whereas the canonical K-K cross-links remained stable, serving as an internal control (Figure 3B).

**Figure 3.**
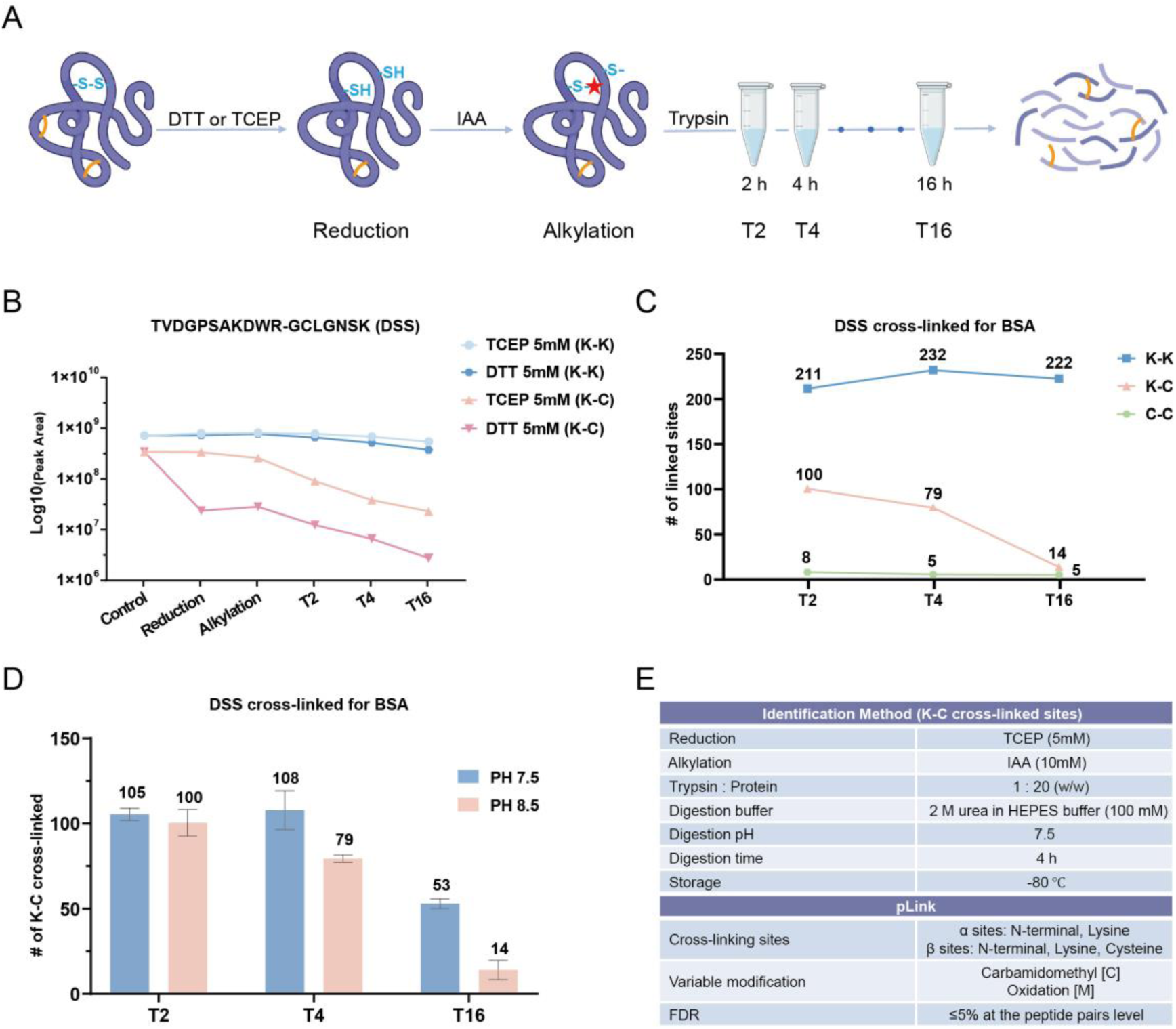
Optimization of XL-MS sample preparation workflows to enhance the stability of lysine-cysteine (K-C) cross-links. (A) Schematic workflow for proteomic sample processing following chemical cross-linking. (B) Comparative stability of K-K and K-C cross-linked peptides during sample preparation. Peak areas of DSS-cross-linked TVDGPSAKDWR and GCLGNSK product, featuring either K-K (blue) or K-C (pink) cross-linked sites, were monitored throughout a urea-based protocol. The impact of reducing agents (5 mM TCEP or DTT) was evaluated over a 16 h time course. (C) Stability of K-K (blue), K-C (pink), and C-C (green) cross-links in BSA as a function of digestion time. Data points represent the total number of identified cross-linked sites using the DSS cross-linker at 2, 4, and 16 h of digestion in 100mM Tris buffer (pH 8.5). (D) Influence of pH and incubation time on K-C cross-link stability. The number of identified K-C sites in DSS-cross-linked BSA was monitored at different time points (2, 4, 16 h) and under different pH conditions (7.5, blue and 8.5, pink) during digestion. (E) Summary of optimized reagents, experimental conditions, and software parameters established for robust K-C cross-link identification. Data are presented from technical replicates (n = 2).

To validate these findings in a complex protein context, we analyzed the data from DSS cross-linked bovine serum albumin (BSA). After a 16-hour tryptic digestion, the number of identified K-C cross-links decreased dramatically from 100 to 14, while the identifications of K-K cross-links remained stable (211 to 222). A shorter, 4-hour digestion was found to be optimal for maximizing the total number of identified cross-links (Figure 3C). Although prolonged digestion progressively reduced K-C cross-links, peptide peak areas reached a stable level after 4 h (Figure S5A), indicating that a 4-hour digestion provides a practical balance between K-C preservation and digestion efficiency. Notably, C-C cross-links were detected at much lower levels, and their numbers did not correlate with digestion time (Figure 3C). Although C-C cross-links are chemically possible, the absence of a systematic trend together with their extremely low frequency suggests that the identified C-C species are dominated by random matches rather than reproducible cross-linking events under our experimental conditions. Therefore, we recommend excluding C-C cross-linked sites from search parameters to minimize the false discovery rate (Figure S5B).

Next, we investigated the effect of pH on K-C cross-link stability, as thioester hydrolysis is known to be base-catalyzed. By comparing digestions of cross-linked BSA at pH 7.5 (50 mM HEPES) versus the more traditional pH 8.5 (100 mM Tris), we found that the near-neutral pH 7.5 buffer effectively mitigated K-C cross-link degradation, particularly during prolonged 16-hour incubations (Figure 3D).

In summary, we identified three key parameters for maximizing the recovery of K-C cross-links: (i) using TCEP for reduction, (ii) maintaining a near-neutral digestion pH of 7.5, and (iii) limiting the digestion time to approximately 4 hours. The detailed experimental conditions and search parameters for the optimized workflow are provided in Figure 3E.

### Application of the Optimized Workflow to Model Proteins

The optimized workflow was validated using four model proteins (BSA, P450, Lactoferrin, and Lysozyme) and three distinct NHS ester-based cross-linkers (DSS, DSSO, and DSBU) (Figure 4A, Figure S8-S11). Significant populations of K-C cross-links were consistently identified alongside canonical K-K sites across all protein scaffolds, demonstrating the general applicability of the method. For BSA, which contains a high number of reactive residues (60 lysines, 35 cysteines), DSS yielded 265 K-K and 142 K-C cross-linked sites. This indicates that incorporating K-C analysis increases the total number of structural constraints by over 50% for this protein. Conversely, in proteins with limited lysine availability, such as Lysozyme (6 lysines, 8 cysteines), K-C cross-linked sites consistently outnumbered K-K identifications across all three reagents; for instance, DSS captured 19 K-C sites compared to only 13 K-K sites. Furthermore, the MS-cleavable cross-linkers DSSO and DSBU exhibited K-C identification efficiencies comparable to the non-cleavable DSS, confirming that acylation of cysteine thiols is a general chemical property of the NHS-ester functional group. These data collectively establish that K-C cross-links are not sporadic artifacts but are reproducible, high-abundance cross-linked sites that significantly expand the landscape of distance constraints available for structural modeling.

**Figure 4.**
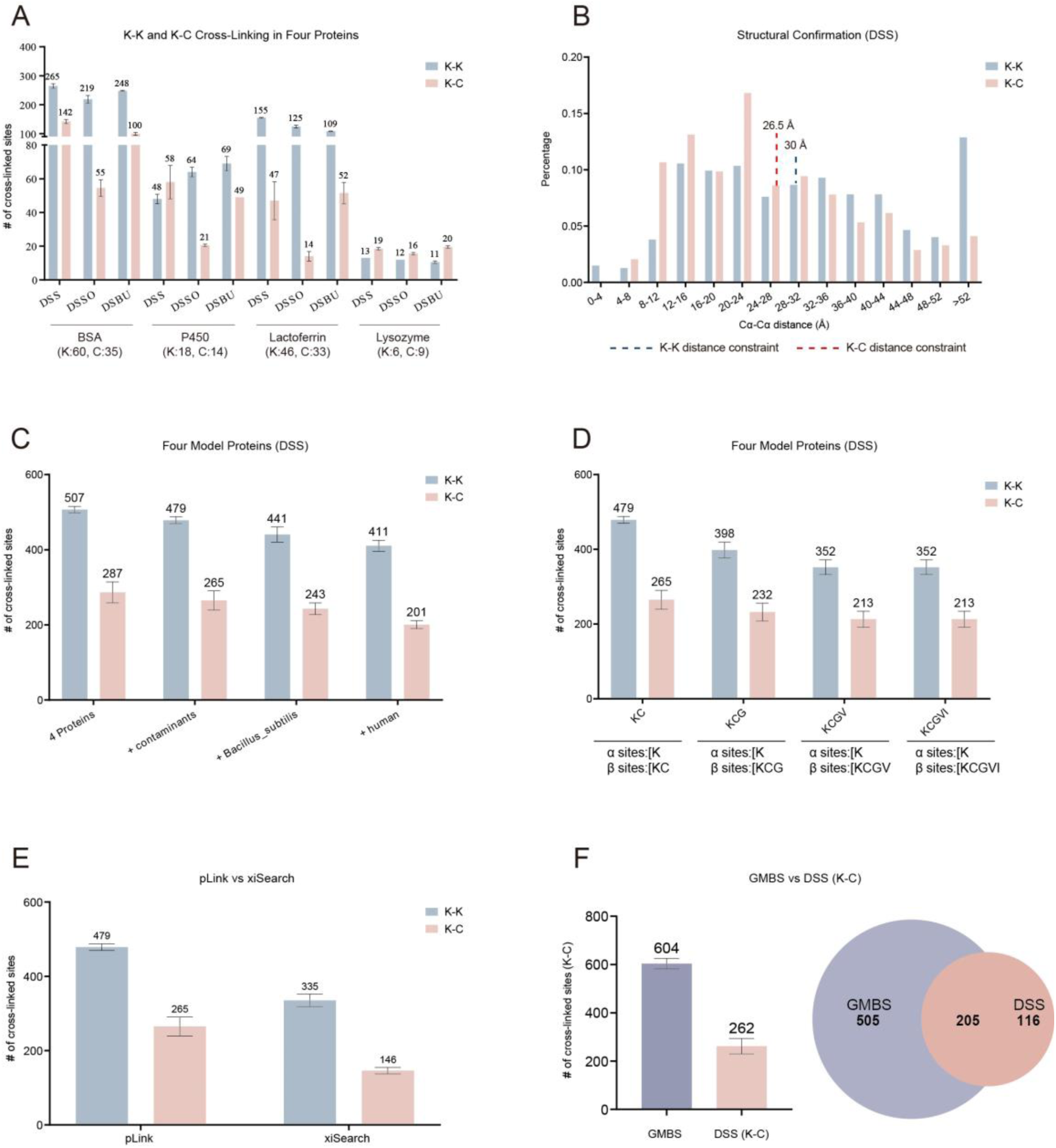
Implementation and multidimensional validation of lysine-cysteine (K-C) cross-link identifications. (A) Identification of K-K (blue) and K-C (pink) cross-linked sites across four proteins with varying Lys/Cys content. Total numbers of identified sites are shown for three distinct reagents (DSS, DSSO, and DSBU) under optimized conditions. The number of Lys/Cys is shown below the figure (B) Structural validation of identified K-K and K-C cross-links. Distribution of Cα-Cα Euclidean distances for K-K (blue) and K-C (pink) cross-linked sites was mapped onto corresponding crystal structures. Dashed lines indicate the theoretical upper-bound constraints (30 Å for K-K and 26.5 Å for K-C). (C) Robustness of K-C cross-link identification against expanding search spaces, ranging from four proteins to the full human proteome. (D) Evaluation of K-C identification via amino acid competition analysis. The number of K-K (blue) and K-C (pink) sites identified when Gly (G), Val (V), and Ile (I) were designated as decoy reactive sites in pLink search parameters. X-axis categories represent the combinations of residues allowed as potential cross-linking sites. (E) Comparison of unique K-K (blue) and K-C (pink) site counts identified by pLink and xiSearch. All searches were conducted using the same raw data with a peptide-pair level FDR ≤ 5%. (F) Distribution and overlap of K-C cross-linked sites identified by GMBS and DSS. The bar plot (left) and intersection (right) compare unique K-C site counts for GMBS (purple) and DSS (pink) across four model proteins. Total site counts are indicated above the bars. The Venn diagram illustrates the overlap between reagent-specific and common K-C cross-linked sites. Data are presented from technical replicates (n = 2).

To assess their structural compatibility, we mapped the identified DSS cross-links onto available crystal structures and calculated the Cα-Cα distances. The distances for both K-K and K-C cross-links were predominantly distributed within the range of 12-40 Å. The distance distribution of K-C cross-links was slightly shorter than that of K-K cross-linked sites, consistent with the theoretical maximum linker spans (26.5 Å vs. 30.0 Å, respectively). Accordingly, 60% of K-C and 51% of K-K cross-links, calculated from the pooled dataset across the four model proteins, satisfied their respective theoretical distance constraints. Similar distributions were observed for DSSO and DSBU (Figure S6A, Figure S7A), confirming that the K-C cross-links are as structurally reliable as canonical K-K cross-links.

The reliability of K-C identifications was further evaluated using two rigorous bioinformatic approaches. First, we performed a search against a large entrapment database (target sequences plus 293 common contaminants and the entire Bacillus subtilis and human proteomes). In this competitive search, the number of K-K identifications decreased from 507 to 411 (a 19% loss), while K-C identifications decreased from 287 to 201 (a 30% loss), indicating comparable sensitivity loss (Figure 4C). Second, we performed a search in which non-reactive amino acids (G, V, I) were included as “entrapment link sites” to capture ambiguous assigned cross-linked sites (Figure 4D). This analysis showed a 26% loss for K-K links (479 to 352) and a 20% loss for K-C links (265 to 213).

Furthermore, to address potential ambiguities arising from search-space expansion and site-localization among alternative nucleophilic residues, we evaluated an extended entrapment strategy by incorporating serine, threonine, and tyrosine (S, T, and Y) as additional candidate cross linked sites. At the cross-link spectrum match (CSM) level, 988 spectra from the original K-C assignments were re-identified with identical peptide backbone sequence pairs. Among these, 84.6% retained their original K-C assignments, with alternative nucleophiles accounting for only a minor fraction of the reassignments (9.4% K-S, 3.9% K-T, 1.6% K-Y, and 0.4% K-K). Similarly, at the peptide-pair level, among 413 peptide-pairs, 85.2% retained their original K-C assignment (Figure S13A-B). These results indicate that K-C assignments remain highly stable even when alternative nucleophilic residues are included in the search. The remaining ∼15% of reassigned spectra most likely reflect increased competition among alternative candidate cross-linking sites within the expanded search space, rather than widespread misidentification of K-C cross-links, supporting the practical reliability of the proposed workflow.

To verify unambiguous site localization for the retained K-C links, we performed a two-part analysis. The sequence distance to the nearest competing residue (K, C, S, T, or Y) averaged 2.6 residues (median: 2) for the retained K-C group (Figure S13C). In contrast, reassigned groups showed significantly shorter distances (1.9 for K-S and 1.0 for K-K), indicating that ambiguity is primarily limited to adjacent sites. In addition, we designed a Local Linked Site Score to quantify the fragment-ion matching within a ±2 fragmentation site window around the linkage, integrating both the number of matched ions and the intensities of the matched peaks:

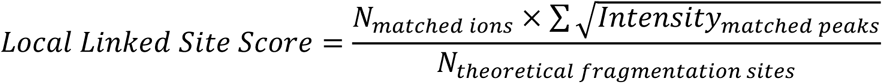

The retained K-C group averaged a high score of 0.9, substantially outperforming reassigned ambiguous clusters (0.4 for K-S and 0.2 for K-K) (Figure S13D), providing additional evidence that the majority of K-C cross-links are supported by strong local fragment-ion evidence and remain reliable under competitive search conditions.

We further strengthened our findings through cross-software validation using pLink and xiSearch (Figure 4E, Figure S6D, S7D). Although pLink reported more identifications overall (479 K-K; 265 K-C) than that from xiSearch (335 K-K; 146 K-C), both platforms consistently detected a significant K-C population, confirming the results are not software-dependent. Finally, we benchmarked our optimized DSS workflow against GMBS, a heterobifunctional “gold-standard” K-C cross-linker. Our method identified 262 K-C cross-links, corresponding to approximately 40% of those identified using GMBS (Figure 4F, Figures S6E and S7E). This result demonstrates that, although DSS does not fully recapitulate the coverage achieved by GMBS, it can recover a substantial subset of K-C cross-links identified by a specialized heterobifunctional reagent. These findings highlight the utility of standard NHS esters, when paired with our optimized protocol, for generating K-C distance constraints.

## Discussion

The findings of this study demonstrate that N-hydroxysuccinimide (NHS) esters—long regarded as lysine-specific reagents—exhibit significant intrinsic reactivity toward cysteine residues. This dual-reactivity transforms conventional NHS-based cross-linking mass spectrometry (XL-MS) into a more versatile tool capable of generating high-quality lysine-cysteine (K-C) cross-linked sites alongside traditional lysine-lysine (K-K) cross-links. The integration of K-C cross-linked sites into standard XL-MS workflows offers several distinct advantages for structural biology. First, by utilizing a single homobifunctional reagent (such as DSS), our approach allows for the simultaneous identification of two distinct types of cross-linking sites. This significantly increases the density of spatial constraints per protein molecule, providing a more granular map of the protein’s tertiary structure and enhancing the resolution of integrative structural modeling. Second, the ability to target cysteine is of particular functional relevance. Cysteine residues are frequently located within the active sites of diverse enzyme classes, including oxidoreductases and transferases. By capturing K-C cross-linked sites in these regions, our method may provide a robust chemical probe to investigate enzyme-substrate interactions and the microenvironments of functional domains that were previously “blind” to lysine-only workflows. Third, and perhaps most importantly from a chemical perspective, this method establishes a reliable pathway for the formation of stable thioester bonds. Given that thioesters are critical intermediates in numerous biochemical processes yet are often transient or labile, our workflow provides a valuable methodology to study the properties, stability, and reactivity of this specific chemical bond within a controlled proteomic context.

Despite these potential advantages, the optimized workflow is tailored for purified proteins and low-complexity samples, and its feasibility at the proteome scale remains to be validated. One primary limitation is the inherent difficulty of identifying cross-links within highly complex proteomes. While sample fractionation or the use of enrichment-tagged cross-linkers can alleviate the signal-to-noise issue, these additional steps inevitably increase sample processing time. This trade-off between proteomic depth and procedural efficiency remains a critical consideration, as the increased complexity of sample preparation may offset the gains in K-C cross-linked site identification. Another major limitation is chemical competition: within complex whole-protein extracts, the vast abundance of lysine residues preferentially and rapidly depletes NHS esters, effectively diluting and suppressing the parallel reactivity toward cysteines. Furthermore, the dependence on the availability of free sulfhydryl groups warrants greater attention (Figure S12A). In proteins stabilized by extensive disulfide networks or located in oxidizing extracellular environments, K-C cross-linking yields are inherently limited by the scarcity of reactive thiols. Crucially, this represents not merely a technical limitation, but a fundamental constraint on the method’s applicability. Consequently, a prior assessment of the protein’s redox state and the quantification of free cysteine content are essential prerequisites to ensure the feasibility of this approach for a given target.

## Supporting Information

The Supporting Information is available free of charge on the ACS Publications website.· Supplementary Methods; experimental design; Figure S1: characterization of cysteine-localized DSS mono-links (pFind); Figure S2: characterization of cysteine-targeted modifications (pFind); Figure S3: MS/MS spectra of synthetic lysine-cysteine cross-linked peptides (pLink); Figure S4: MS/MS spectra of lysine-cysteine cross-linked peptides (public datasets, pLink); Figure S5: evaluation of digestion efficiency and cross-linking search parameters; Figure S6: multidimensional validation of K-C cross-link identifications (DSSO); Figure S7: multidimensional validation of K-C cross-link identifications (DSBU); Figure S8: MS/MS spectra of K-C cross-linked peptides (BSA, DSS); Figure S9: MS/MS spectra of K-C cross-linked peptides (Lactoferrin, DSS); Figure S10: MS/MS spectra of K-C cross-linked peptides (Lysozyme, DSS); Figure S11: MS/MS spectra of K-C cross-linked peptides (P450, DSS); Figure S12: detection of free sulfhydryl groups and crosslinking of new versus old BSA; Figure S13: Validation of Cysteine-Specific Assignments against Alternative Nucleophilic Ambiguities.

## Acknowledgements

This work was supported by the Innovative Drug Research and Development National Science and Technology Major Project (No. 2025ZD1804100). We thank Dr. Yong Cao at the National Institute of Biological Sciences, Beijing, for his support of this work.

## Data Availability Statement

The mass spectrometry proteomics data have been deposited to the ProteomeXchange Consortium (https://proteomecentral.proteomexchange.org) via the iProX partner repository with the dataset identifier PXD075905.

The pFind-3.2.2 and pLink-3.0.17 are available at: https://pfind.ict.ac.cn

## Author Contributions

G.-C.S. and X.-M.Z. devised the project. G.-C.S. and X.-F.Y. designed the experiments in this study. X.-F.Y. and A.-C.H. performed wet-lab experiments. X.-F.Y., A.-C.H., P.-Z.M. and Q.S. performed data analysis. Q.S. performed the schematic diagram preparation. P.-Z.M. and Y.-J.Q. provided mass spectrometry technical support. G.-C.S. and X.-F.Y. wrote the manuscript.

## Note

The authors declare no competing financial interest. This research did not involve human or animal participants.

