## Supplemental for "Enhancing the Identification of NHS Ester-Mediated Lysine-Cysteine Cross-Linking via Reduced Trypsin Digestion Time"

**Methods**

**Reaction of NEM with BSA**

An equal mass of NEM (30 μg, final concentration 8 mM) was added to BSA, followed by incubation at 600 rpm for 1 h. The reaction was then terminated by precipitation with pre-chilled acetone at -20°C for ≥ 2 h. The protein pellet was resolubilized in 8 M urea. Reduction and alkylation were performed using TCEP(5mM) and IAA(10mM), respectively. The solution was subsequently diluted to reduce the urea concentration to 2 M, and digested with trypsin at a 1:20 (w/w) enzyme-to-substrate ratio for 4 h. Finally, the peptides were desalted prior to LC-MS/MS analysis.

**Label Transfer Assay**

A synthetic peptide bearing an N-terminal Boc protecting group (Boc-YLECSALTQR) was conjugated to NHS-Biotin. Following acid-mediated deprotection with 95% trifluoroacetic acid (TFA), the peptide was reconstituted in 50 mM HEPES (pH 7.5). Temporal sampling was conducted over a 16-h kinetic window (0, 0.5, 2, 4, and 16 h) for mass spectrometric analysis. As a negative control, a parallel reaction was performed using the intact, non-deprotected peptide without TFA treatment.

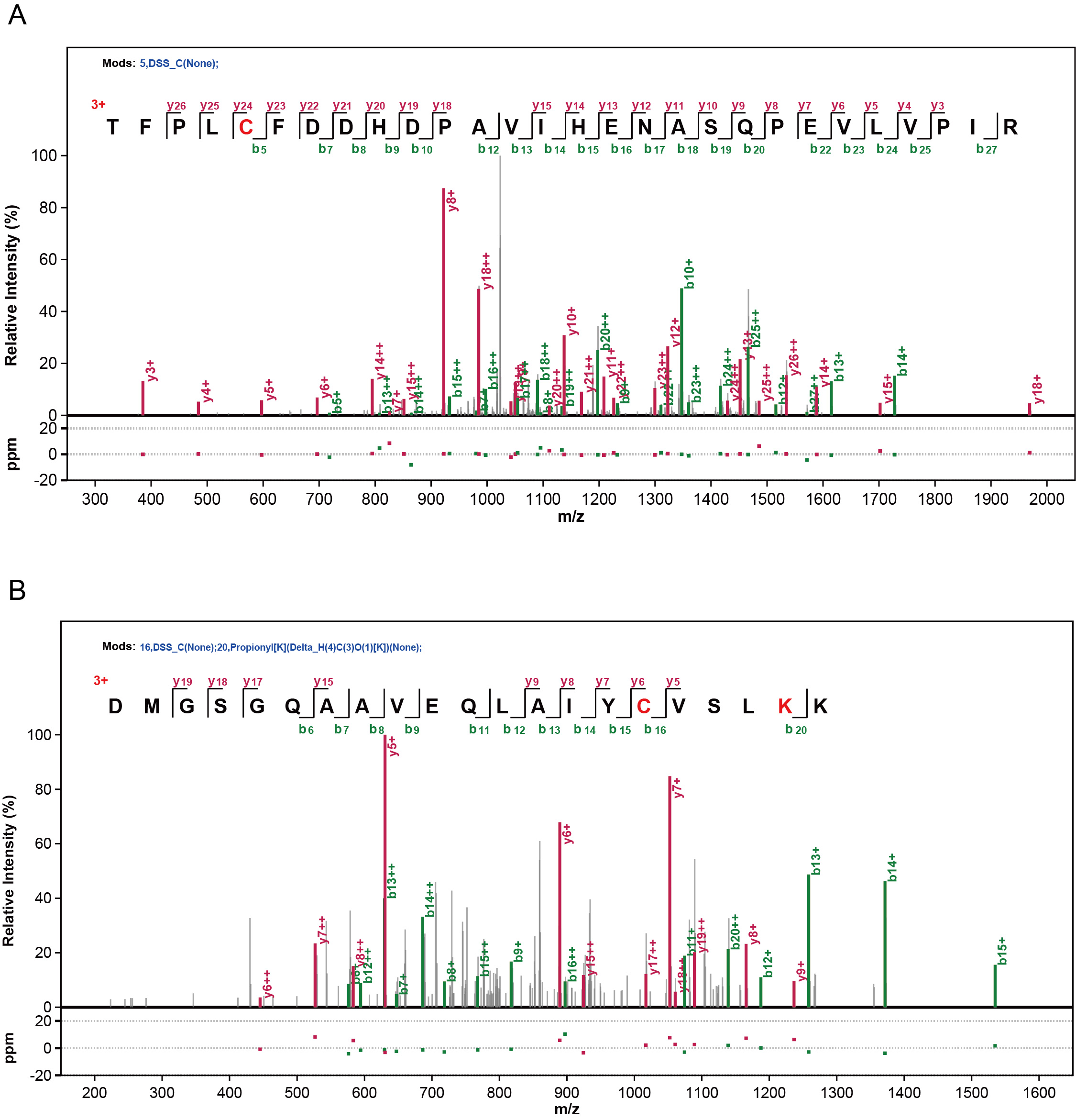

Supplementary Figure 1: Characterization of cysteine-localized DSS mono-links by tandem mass spectrometry (pFind).

Annotated MS/MS spectra of peptides with a DSS mono-link at a cysteine residue. (A) Peptide sequence, TFPLCFDDHDPAVIHENASQPEVLVPIR, z=3+, from Dataset PXD054757, with the DSS mono-link at Cys-5. (B) Peptide sequence, DMGSGQAAVEQLAIYCVSLKK, z=3+, from Dataset PXD058941, with the DSS mono-link at Cys-16. Matched b-ions and y-ions are colored green and red, respectively.

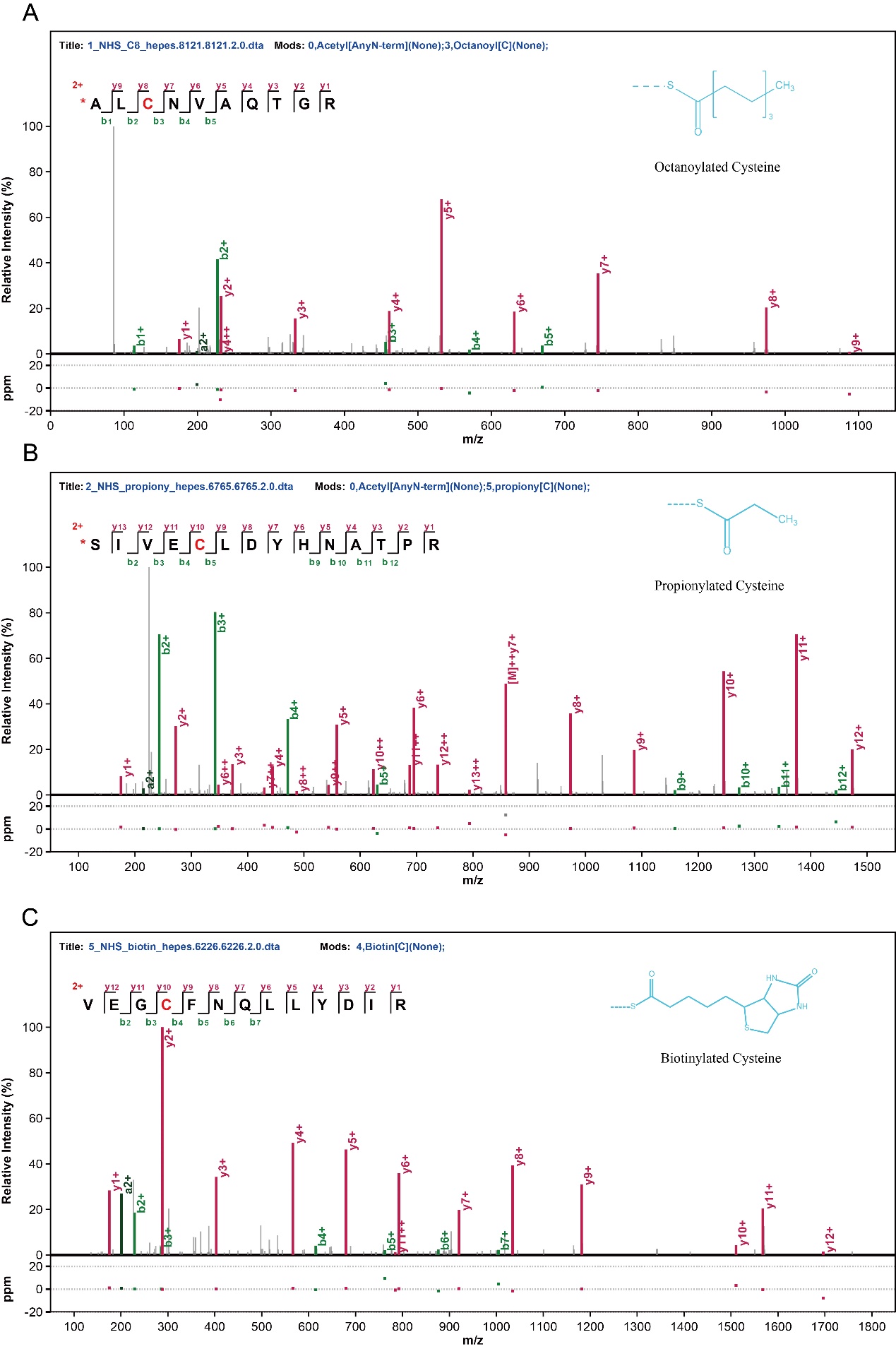

Supplementary Figure 2: Characterization of cysteine-targeted modifications by tandem mass spectrometry (pFind).

Annotated MS/MS spectra show fragment ion matches for: (A) octanoylated cysteine within the peptide, Ac-ALCNVAQTGR, z=2+, (B) propionylated cysteine within the peptide, Ac-SIVECLDYHNATPR, z=2+, and (C) biotinylated cysteine within the peptide, VEGCFNQLLYDIR, z=2+. Matched b-ions (green) and y-ions (red) are annotated for each sequence. The specific modification sites were assigned using pFind. Experimental conditions for all samples: 50 mM HEPES (pH 7.5).

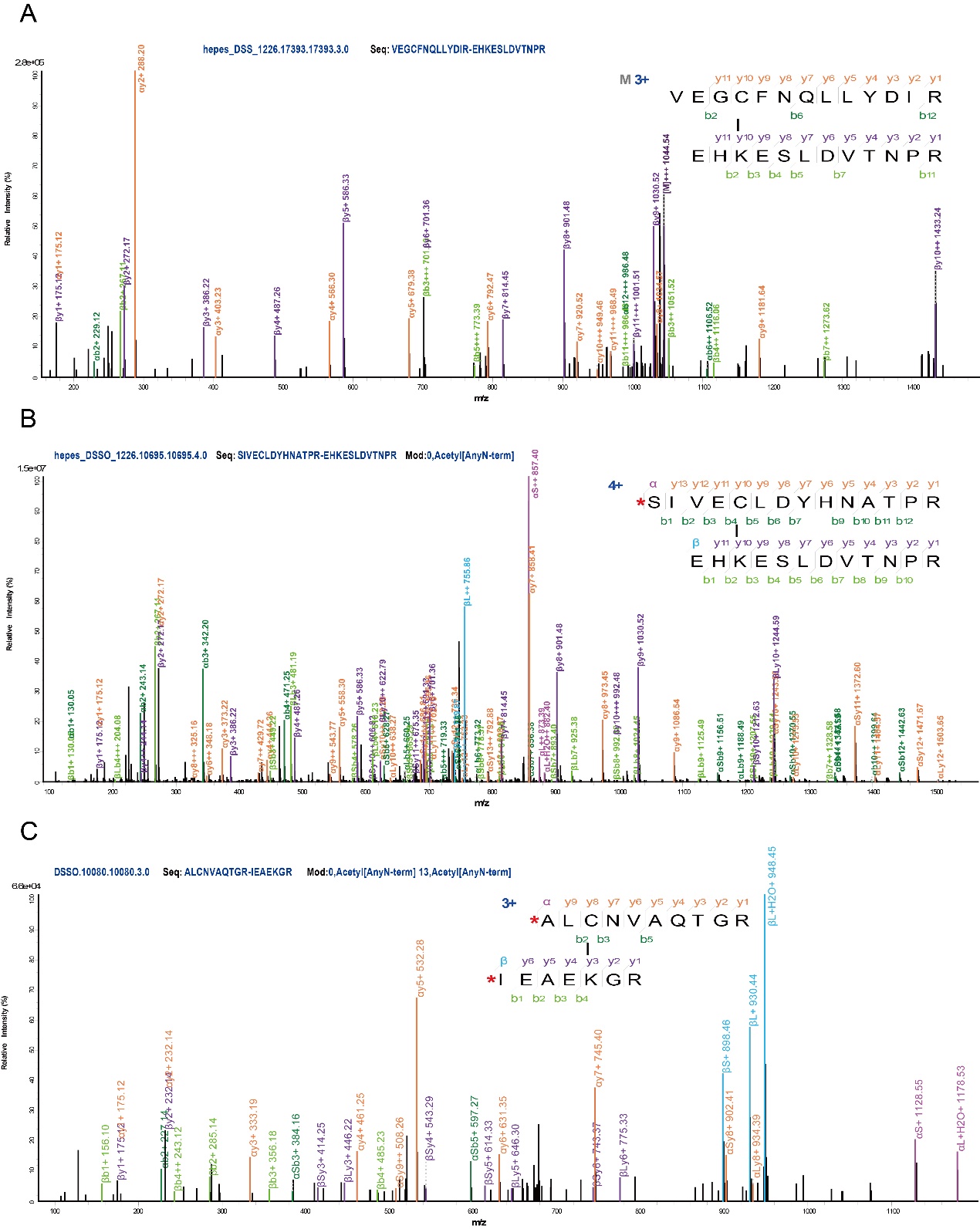

Supplementary Figure 3: MS/MS spectra of the lysine-cysteine cross-linked peptides (synthetic peptide, pLink).

(A) VEGCFNQLLYDIR-EHKESLDVTNPR, m/z=1044.521, z=3+. (B) ac-SIVECLDYHNATPR-EHKESLDVTNPR, m/z=811.127, z=4+. (C) ac-ALCNVAQTGR-ac-IEAEKGR, m/z=692.666, z=3+.

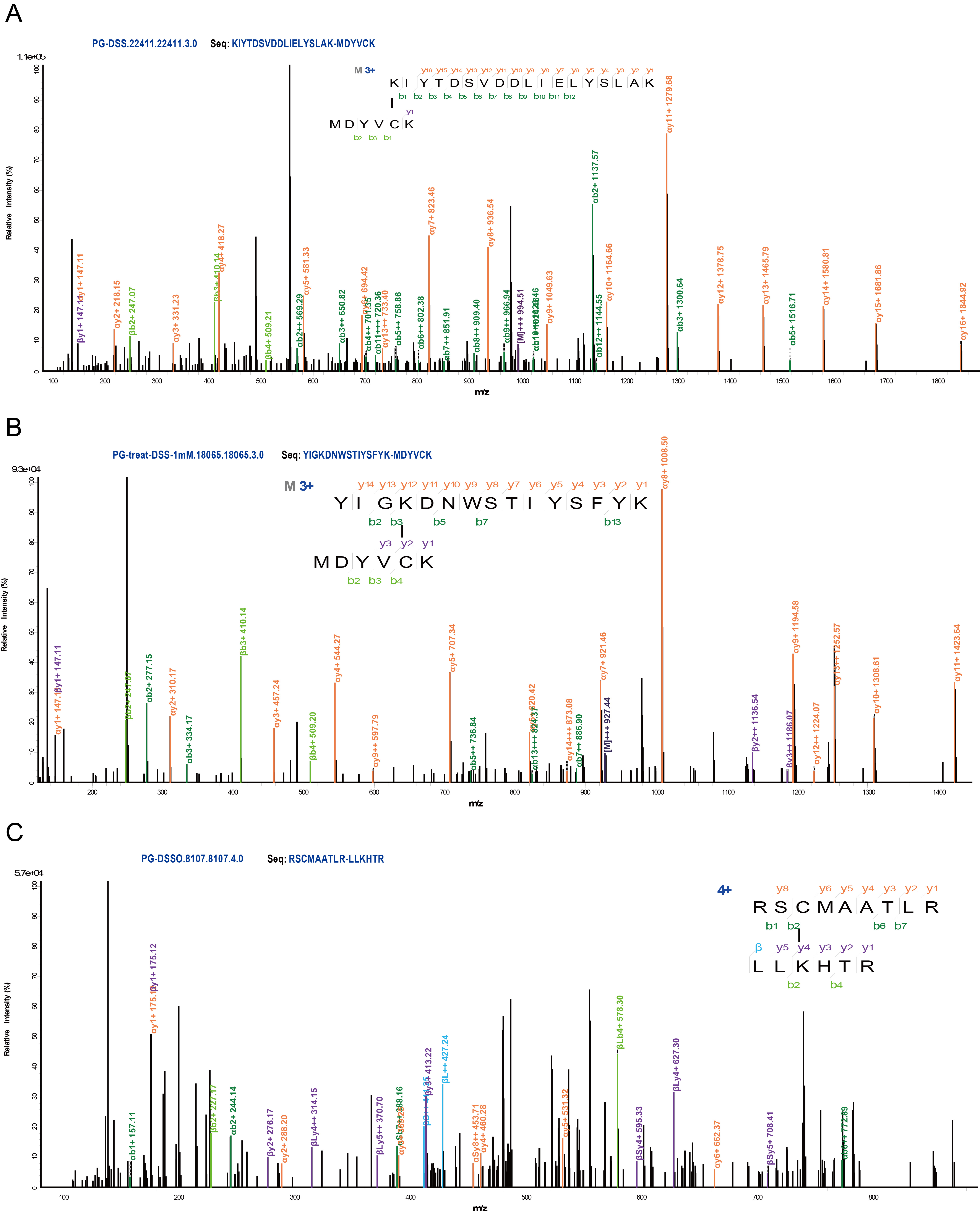

Supplementary Figure 4: MS/MS spectra of the lysine-cysteine cross-linked peptides. (Public datasets, pLink)

The cross-linked peptide (A) KIYTDSVDDLIELYSLAK-MDYVCK, z=3+, and (B) YIGKDNWSTIYSFYK-MDYVCK, z=3+, were identified from the PG-complex following DSS-mediated cross-linking. (C) Cross-linked peptide RSCMAATLR-LLKHTR, z=4+ was identified from the PG-complex following DSSO-mediated cross-linking.

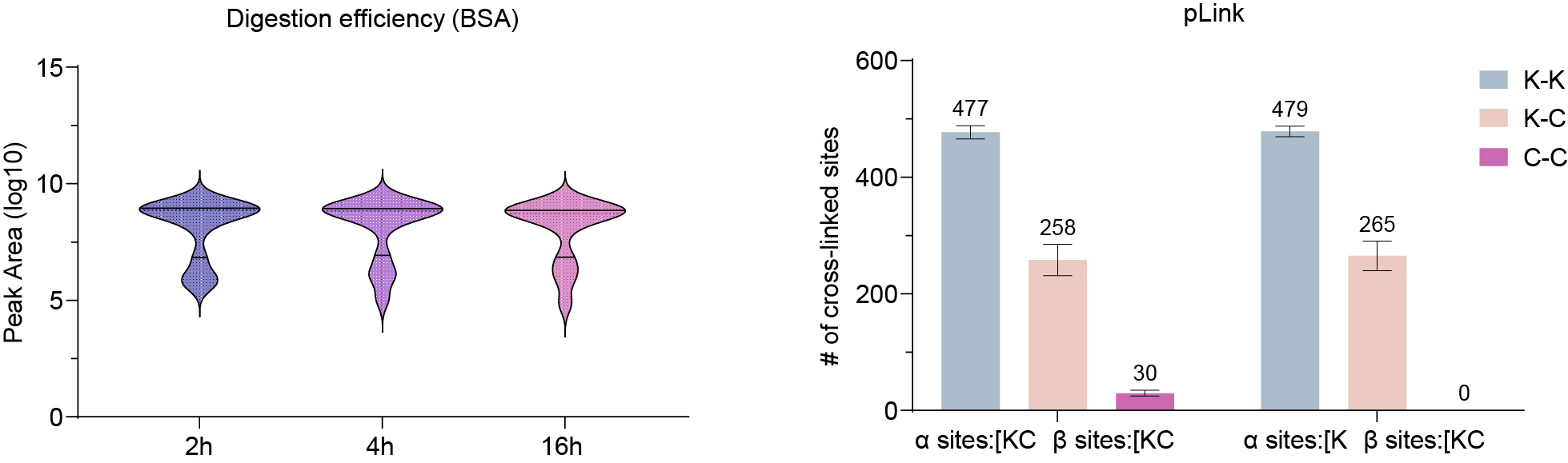

Supplementary Figure 5. Evaluation of digestion efficiency and cross-linking search parameters.

(A) Distribution of peptide peak areas during BSA digestion. Violin plots illustrate the log10 transformed peak areas of identified peptides (n=15) at 2 h, 4 h, and 16 h incubation intervals. Within each violin, the dashed and solid horizontal lines indicate the median and the quartiles, respectively. (B) Distribution of cross-linked sites identified under varying pLink search parameters. The number of unique K-K (blue), K-C (pink), and C-C (red) cross-linked sites is displayed for two distinct search configurations: one including both Lys (K) and Cys (C) as potential α and β sites (α sites:[KC, β sites:[KC), and another restricting α sites to Lys only (α sites:[K, β sites:[KC). Absolute counts of identified unique sites are labeled above each bar. Data are presented from technical replicates (n = 2).

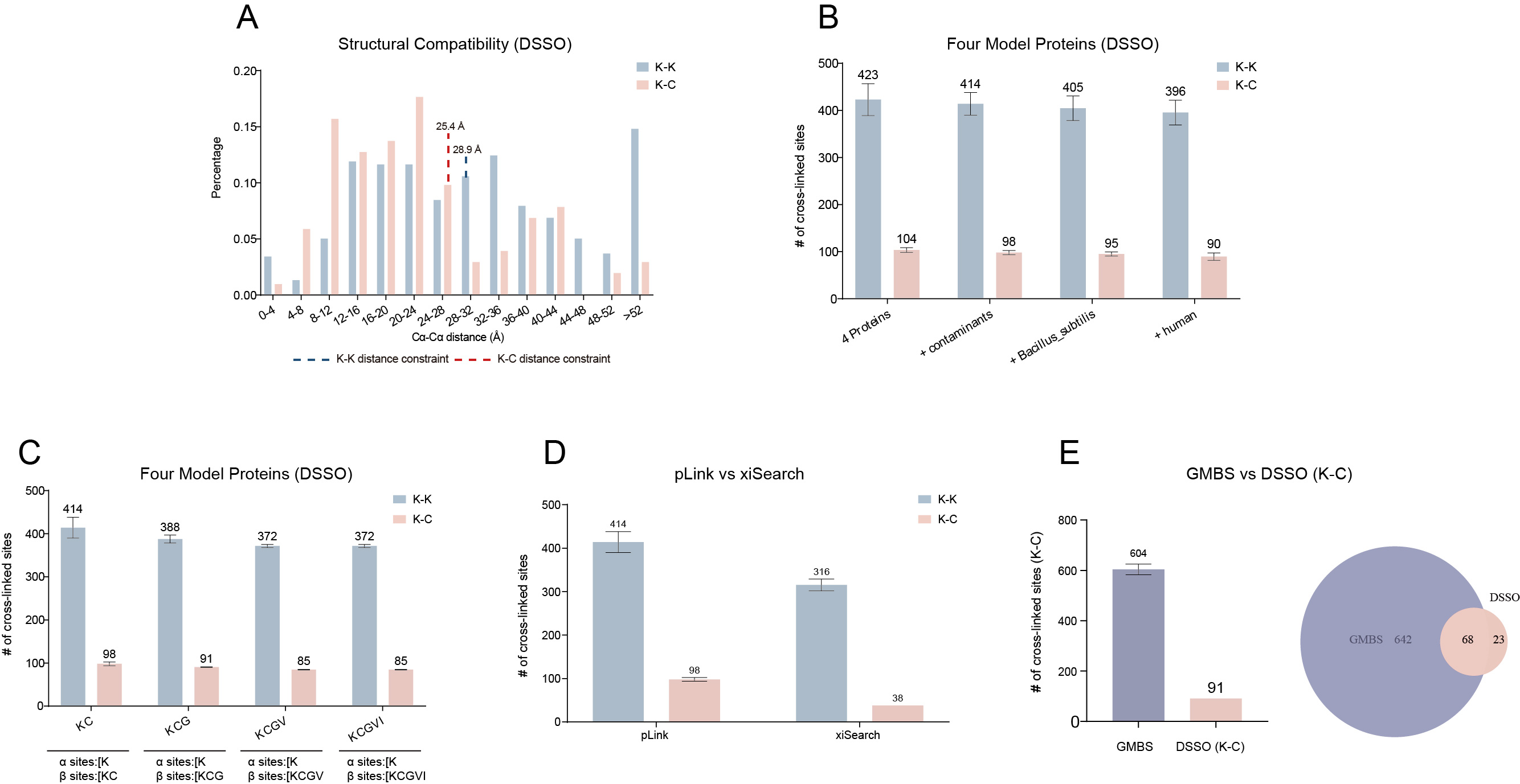

Supplementary Figure 6: Multidimensional validation of lysine-cysteine (K-C) cross-link identifications (DSSO).

(A) Structural validation of identified K-K and K-C cross-links. Distribution of Cα-Cα Euclidean distances for K-K (blue) and K-C (pink) linkages was mapped onto corresponding crystal structures. Dashed lines indicate the theoretical upper-bound constraints (28.9 Å for K-K and 25.4 Å for K-C). (B) Robustness of K-C cross-link identification against expanding search spaces, ranging from four proteins to the full human proteome. (C) Evaluation of K-C identification via amino acid competition analysis. The number of K-K (blue) and K-C (pink) sites identified when Gly (G), Val (V), and Ile (I) were designated as decoy reactive sites in pLink search parameters. X-axis categories represent the combinations of residues allowed as potential cross-linking sites. (D) Comparison of unique K-K (blue) and K-C (pink) site counts identified by pLink and xiSearch. All searches were conducted using the same raw data with a peptide-pair level FDR ≤ 5%. (E) Distribution and overlap of K-C cross-linked sites identified by GMBS and DSSO. The bar plot (left) and intersection (right) compare unique K-C site counts for GMBS (purple) and DSSO (pink) across four model proteins. Total site counts are indicated above the bars. The Venn diagram illustrates the overlap between reagent-specific and common K-C linkages. Data are presented from technical replicates (n = 2).

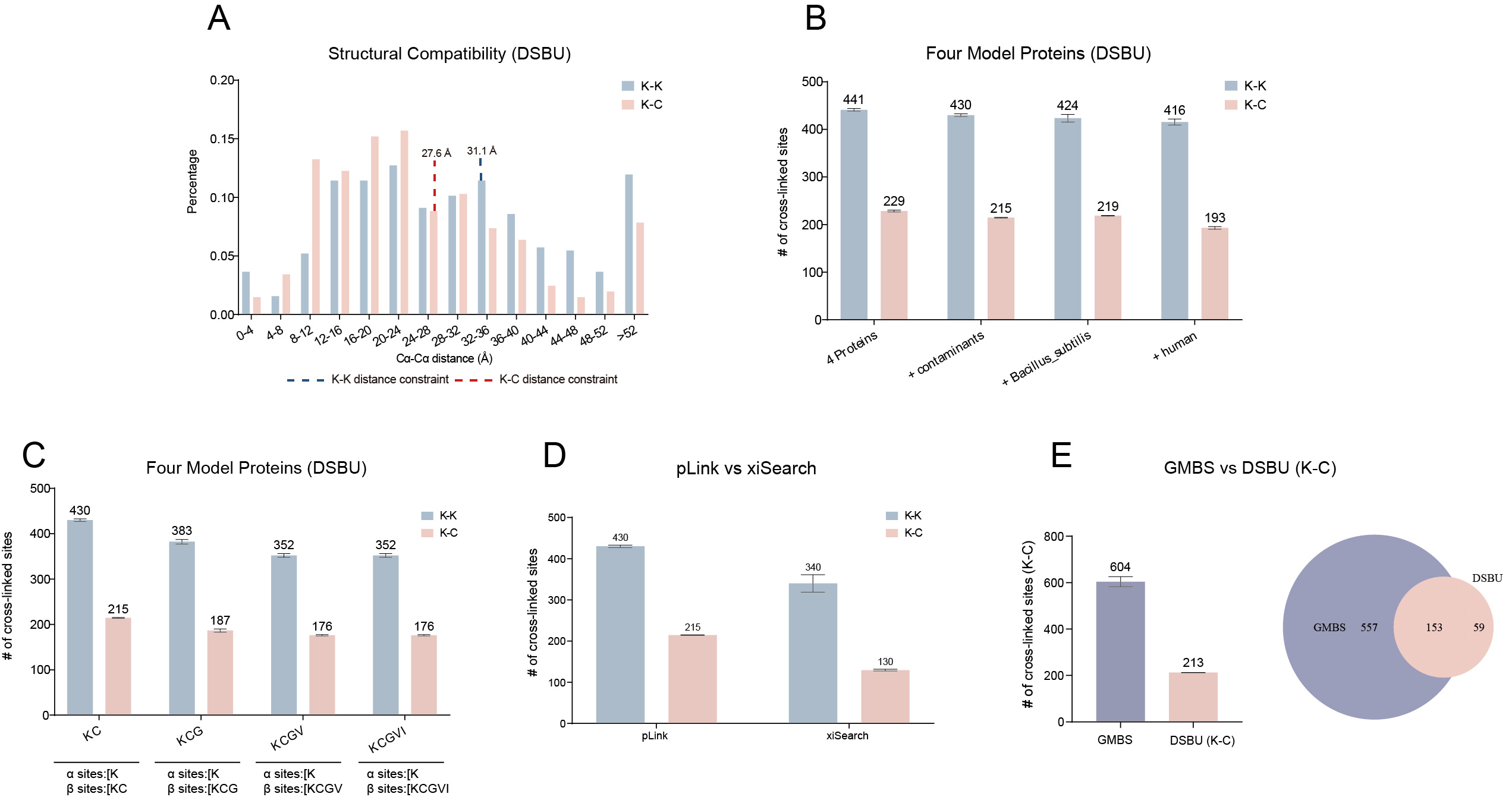

Supplementary Figure 7: Multidimensional validation of lysine-cysteine (K-C) cross-link identifications (DSBU).

(A) Structural validation of identified K-K and K-C cross-links. Distribution of Cα-Cα Euclidean distances for K-K (blue) and K-C (pink) linkages was mapped onto corresponding crystal structures. Dashed lines indicate the theoretical upper-bound constraints (31.1 Å for K-K and 27.6 Å for K-C). (B) Robustness of K-C cross-link identification against expanding search spaces, ranging from four proteins to the full human proteome. (C) Evaluation of K-C identification via amino acid competition analysis. The number of K-K (blue) and K-C (pink) sites identified when Gly (G), Val (V), and Ile (I) were designated as decoy reactive sites in pLink search parameters. X-axis categories represent the combinations of residues allowed as potential cross-linking sites. (D) Comparison of unique K-K (blue) and K-C (pink) site counts identified by pLink and xiSearch. All searches were conducted using the same raw data with a peptide-pair level FDR ≤ 5%. (E) Distribution and overlap of K-C cross-linked sites identified by GMBS and DSBU. The bar plot (left) and intersection (right) compare unique K-C site counts for GMBS (purple) and DSBU (pink) across four model proteins. Total site counts are indicated above the bars. The Venn diagram illustrates the overlap between reagent-specific and common K-C linkages. Data are presented from technical replicates (n = 2).

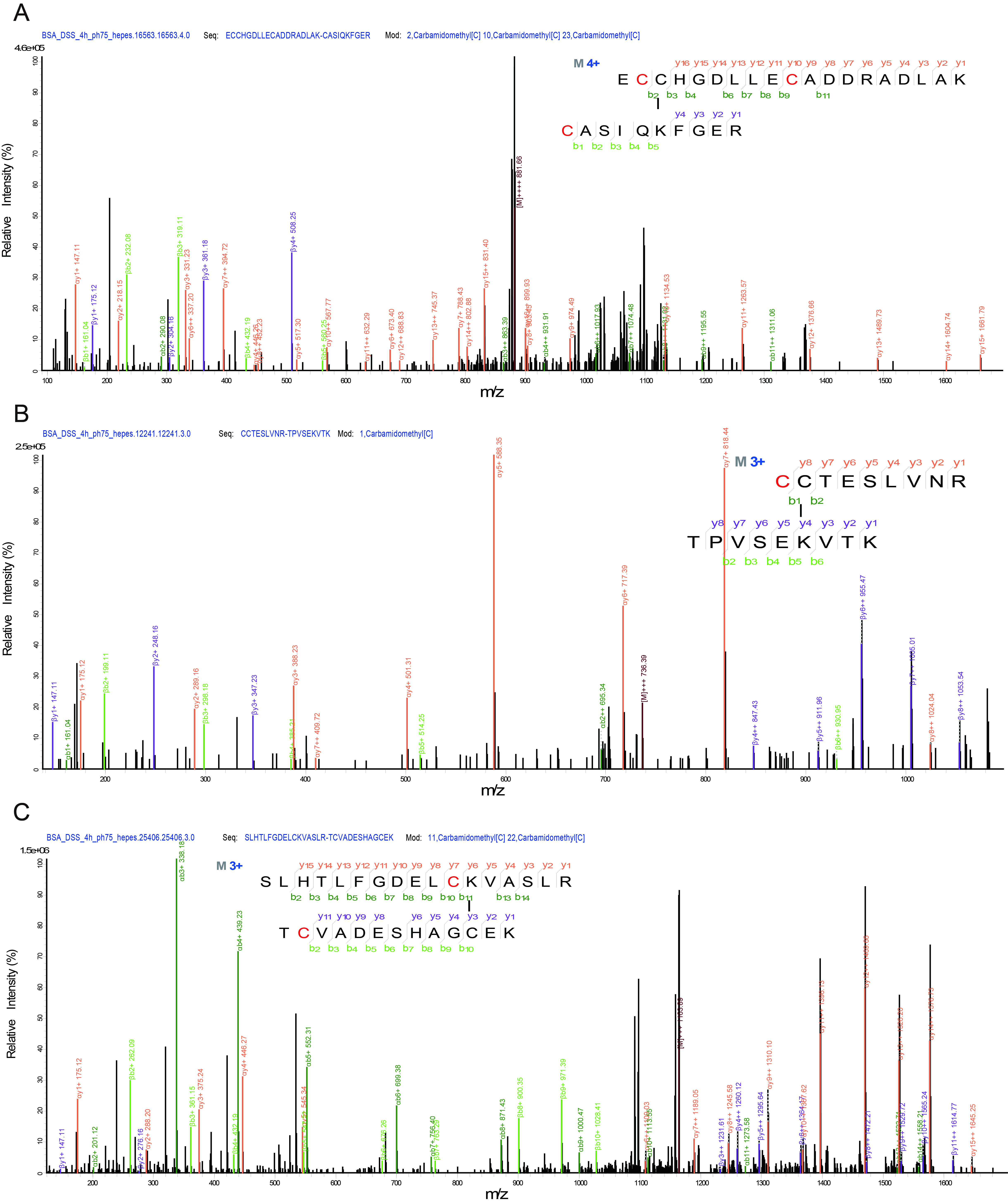

Supplementary Figure 8: MS/MS spectra of the lysine-cysteine cross-linked peptides. (BSA, DSS)

(A) ECCHGDLLECADDRADLAK-CASIQKFGER, MH+= 3523.57, z= 4+. (B) CCTESLVNR-TPVSEKVTK, MH+= 2207.10, z= 3+. (C) SLHTLFGDELCKVASLR-TCVADESHAGCEK, MH+= 3489.64, z= 3+.

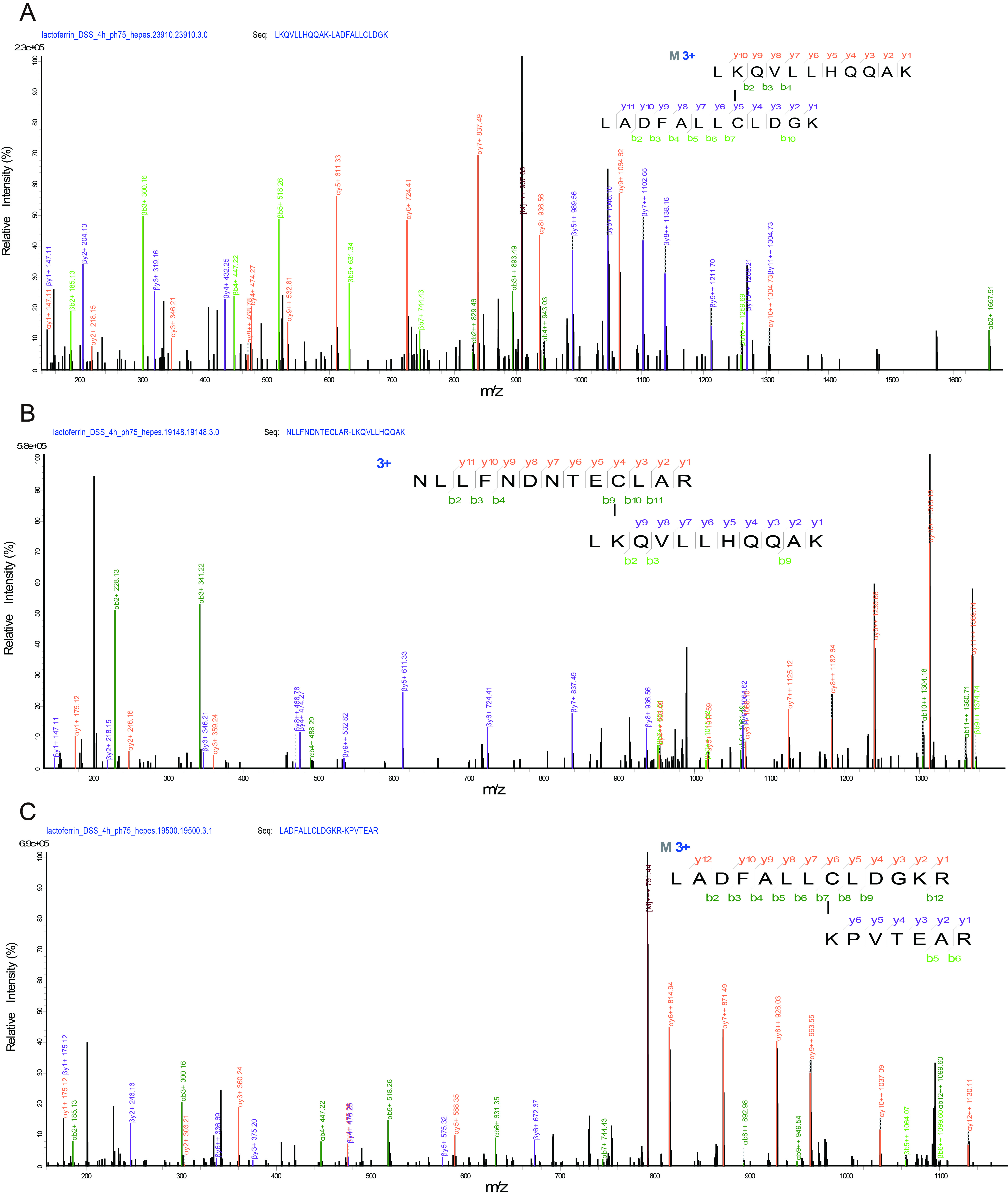

Supplementary Figure 9: MS/MS spectra of the lysine-cysteine cross-linked peptides. (Lactoferrin, DSS)

(A) LKQVLLHQQAK-LADFALLCLDGK, MH+= 2721.53, z= 3+. (B) NLLFNDNTECLAR-LKQVLLHQQAK, MH+= 2965.59, z= 3+. (C) LADFALLCLDGKR-KPVTEAR, MH+= 2372.30, z= 3+.

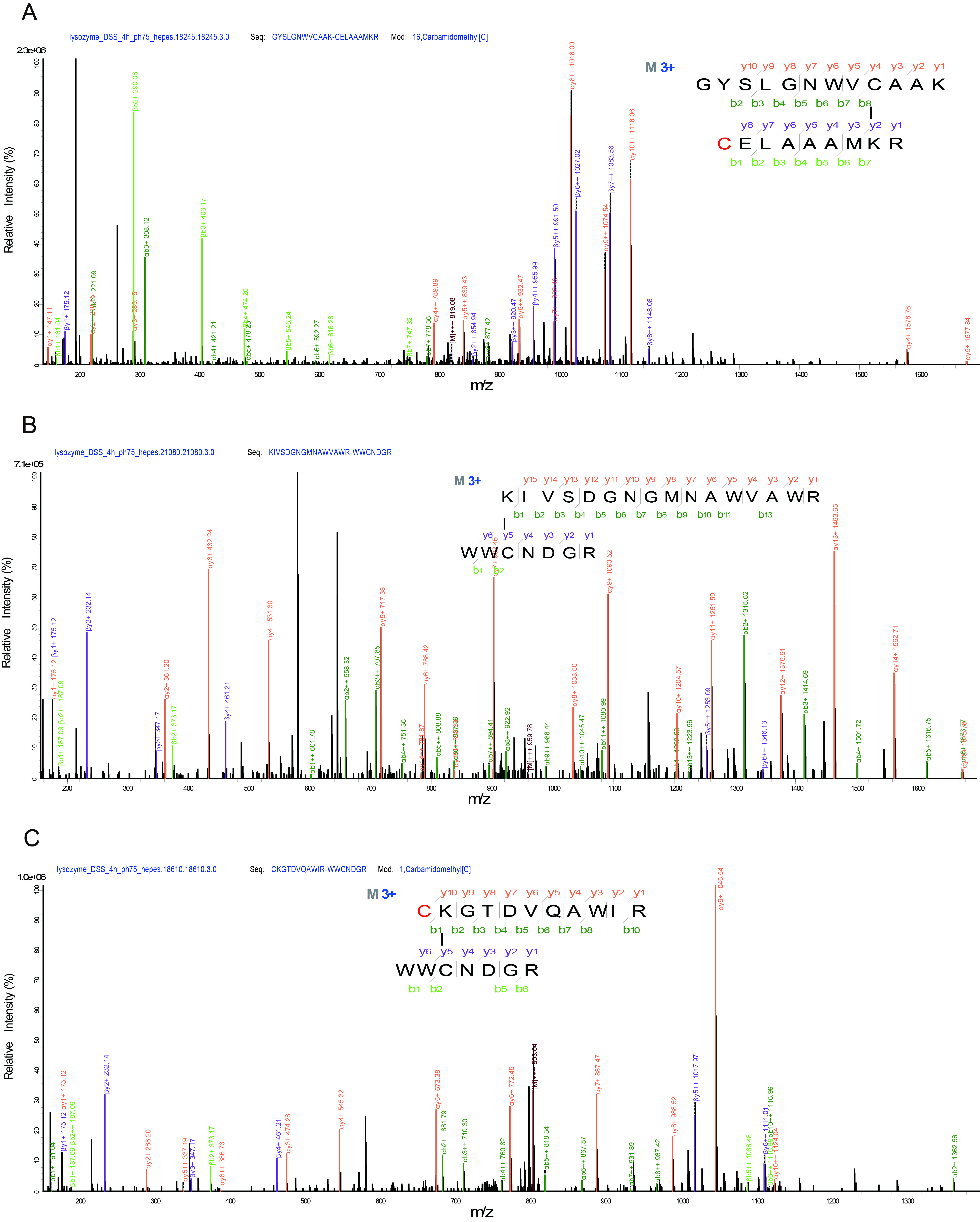

Supplementary Figure 10: MS/MS spectra of the lysine-cysteine cross-linked peptides. (Lysozyme, DSS)

(A) GYSLGNWVCAAK-CELAAAMKR, MH+= 2455.19, z= 3+. (B) KIVSDGNGMNAWVAWR-WWCNDGR, MH+= 2877.33, z= 3+. (C) CKGTDVQAWIR-WWCNDGR, MH+= 2407.10, z= 3+.

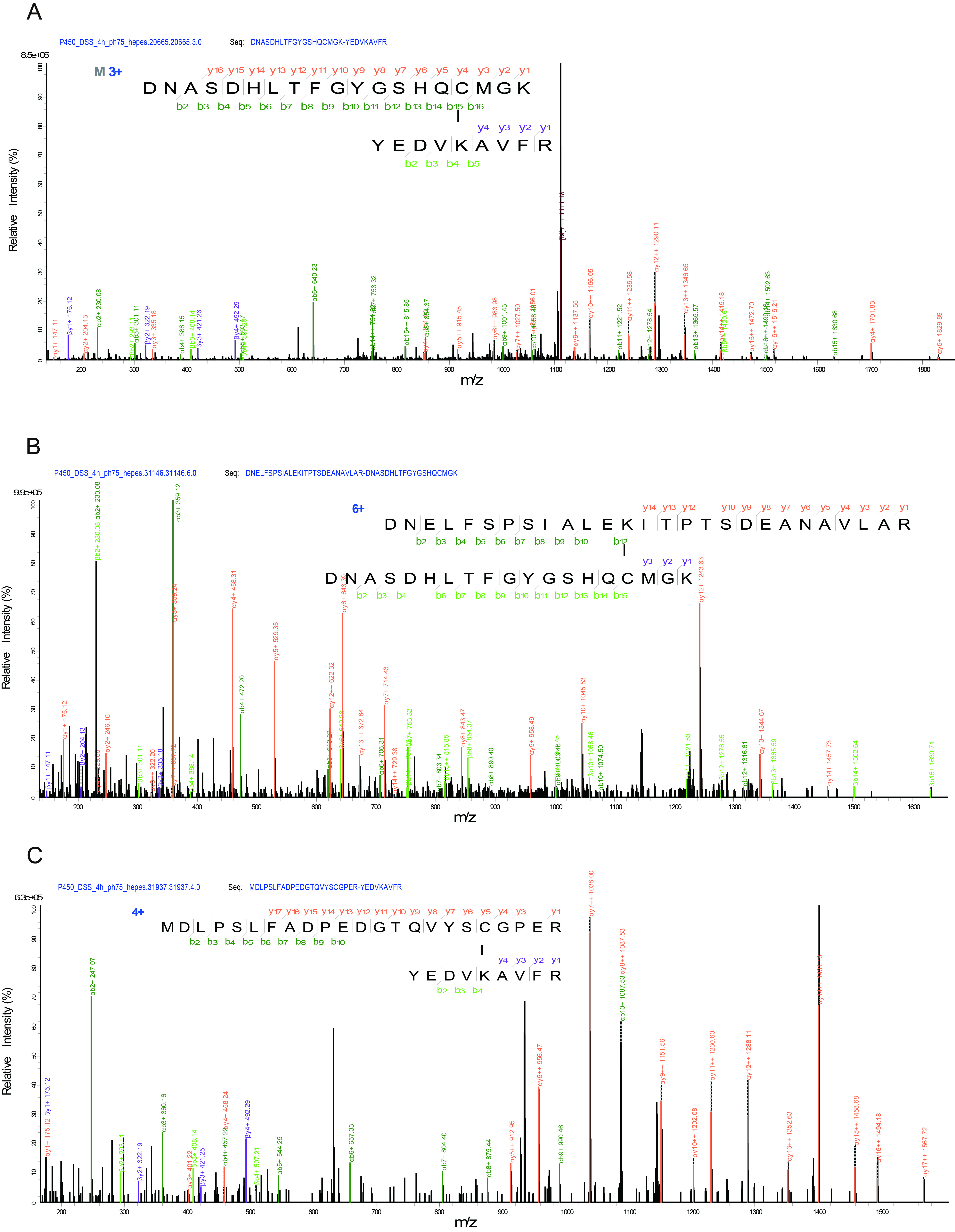

Supplementary Figure 11: MS/MS spectra of the lysine-cysteine cross-linked peptides. (P450, DSS)

(A) DNASDHLTFGYGSHQCMGK-YEDVKAVFR, MH+= 3331.51, z= 3+. (B) DNELFSPSIALEKITPTSDEANAVLAR-DNASDHLTFGYGSHQCMGK, MH+= 5106.41, z= 6+. (C) MDLPSLFADPEDGTQVYSCGPER-YEDVKAVFR, MH+= 3790.76, z= 4+.

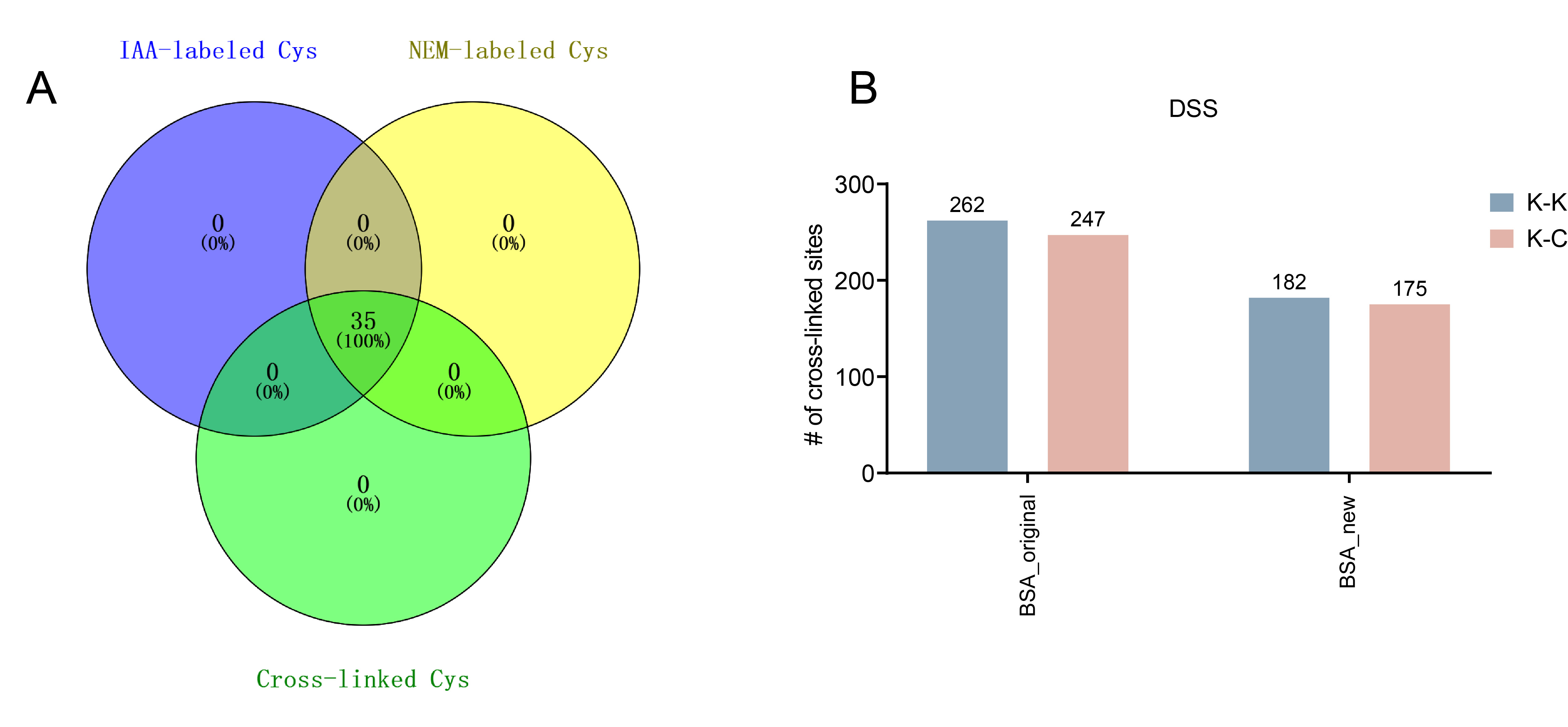

Supplementary Figure 12: Detection of Free Sulfhydryl Groups and Crosslinking of New versus Old BSA.

(A) Carbamidomethyl (derived from IAA modification) denotes Cys residues originally involved in disulfide bonds; N-ethylmaleimide (derived from NEM modification) denotes free thiol-containing Cys; “Crosslinked” denotes Cys residues engaged in crosslinking. (B) DSS was used to simultaneously cross-link new and old BSA. (K-K, blue ; K-C, pink).

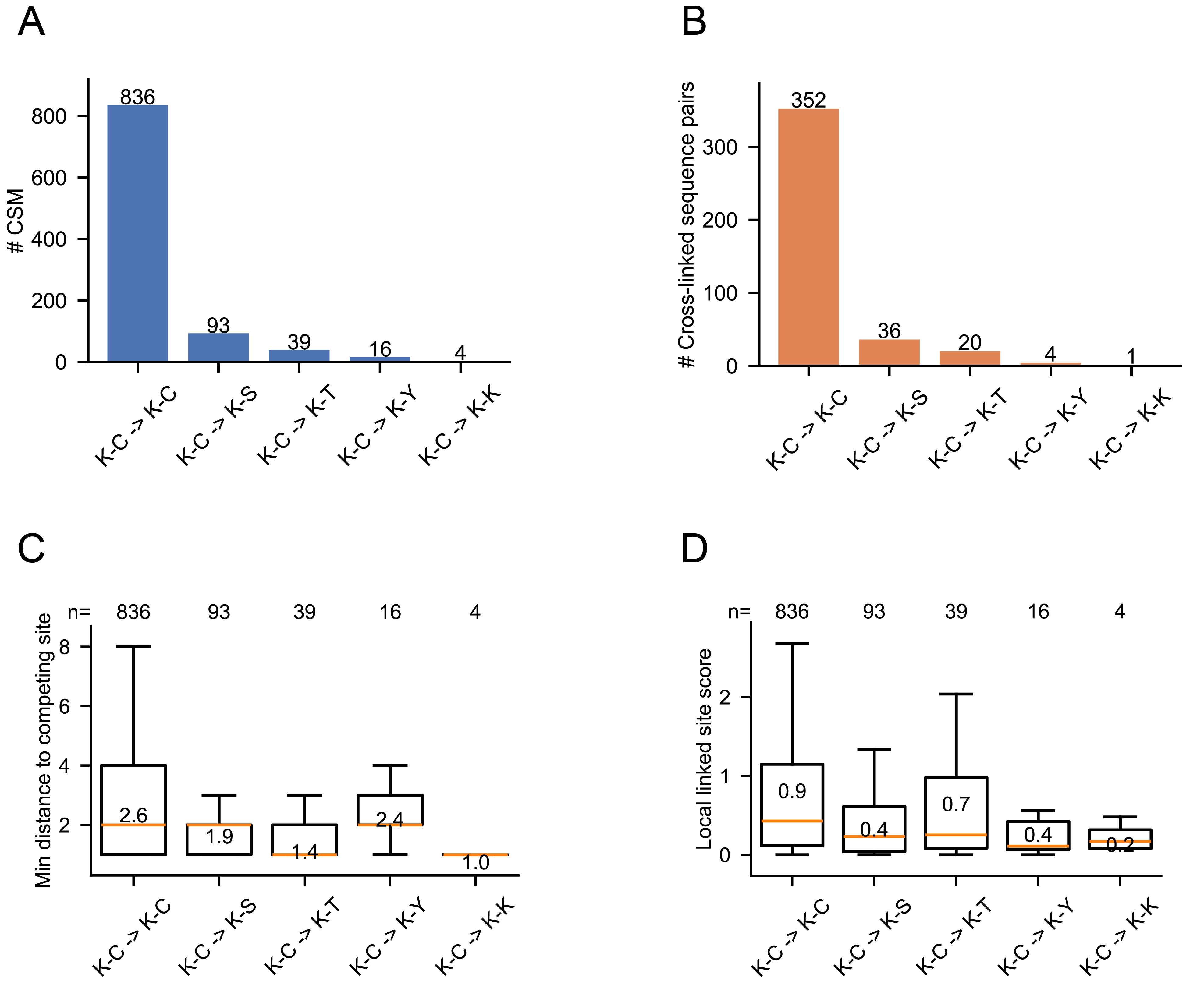

Supplementary Figure 13: Validation of Cysteine-Specific Assignments against Alternative Nucleophilic Ambiguities.

(A-B) Redistribution of originally assigned K-C cross-links following strict competitive re-searching with expanded reactive residue settings (α: [K; β: [KCSTY), analyzed at both the cross-link spectrum match (CSM) level (A) and cross-linked peptide pair level (B). The number of identifications for each category is indicated above the corresponding bars. (C) Sequence distance analysis of cysteine cross-linking sites relative to competing nucleophilic residues in K-C cross-links. Boxplots show the minimum sequence distance between the assigned cysteine cross-linking site and the nearest alternative reactive residue (K, C, S, T, or Y) on the same peptide. Mean values are indicated, and medians are shown as orange lines. (D) Distribution of the Local Linked Site Score across different cross-link categories. Boxplots represent a localized fragmentation-based score calculated within a ±2 fragmentation site window surrounding the assigned cross-linking position.

Supplementary Figure 14: Re-analyzed crosslinking site counts in four proteins.

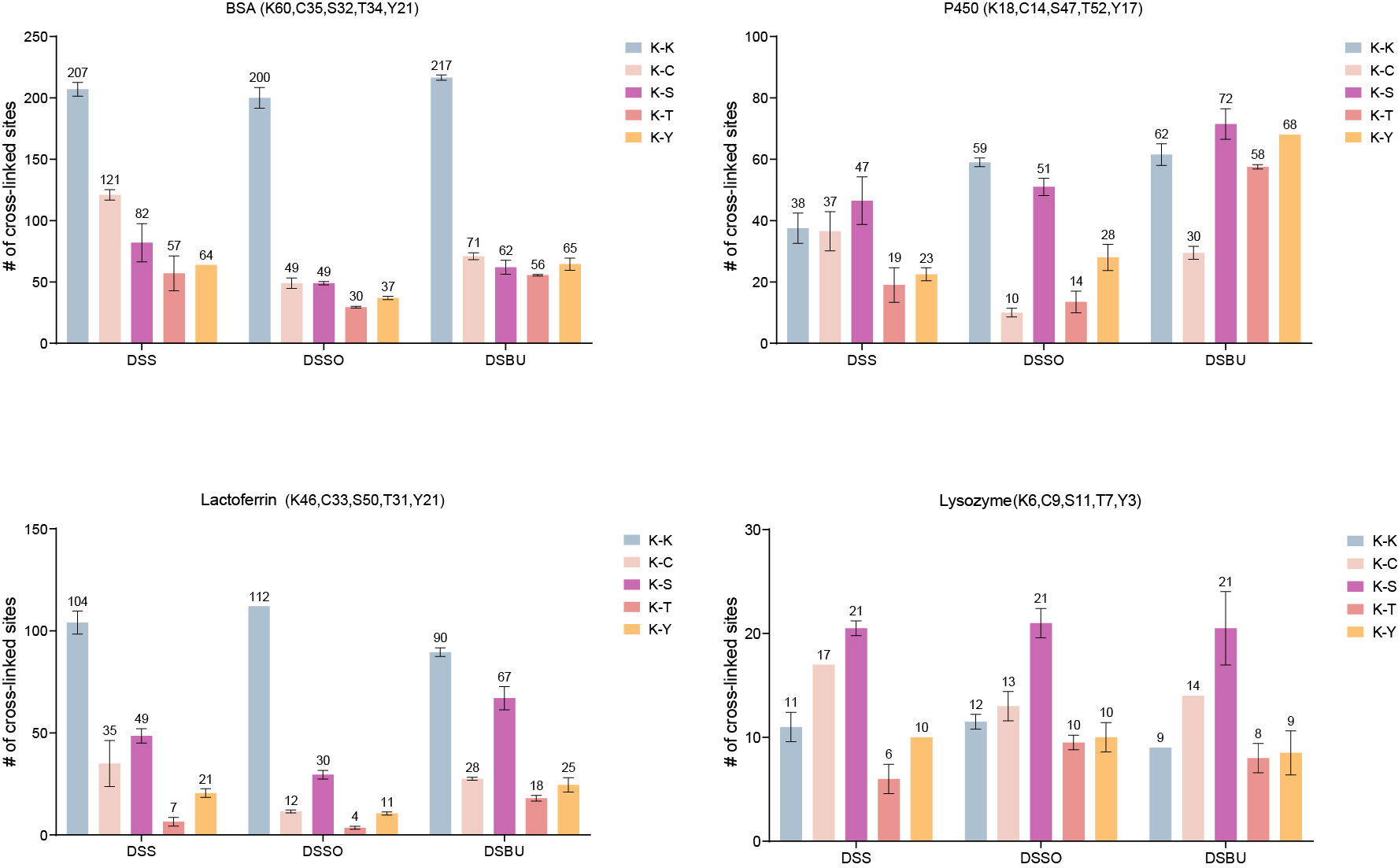

α sites: [K, and β sites: [K, C, S, T, Y . Following process optimization, data were re-analyzed to quantify the potential crosslinking sites for each of the four proteins. Numbers in parentheses indicate the count of the specified amino acid residue within each protein sequence (e.g., BSA contains 60 K residues).
